# The metabolite-controlled ubiquitin conjugase Ubc8 promotes mitochondrial protein import by enhancing assembly of the TOM complex

**DOI:** 10.1101/2022.04.26.489513

**Authors:** Saskia Rödl, Fabian den Brave, Markus Räschle, Svenja Lenhard, Carina Groh, Hanna Becker, Jannik Zimmermann, Bruce Morgan, Elke Richling, Thomas Becker, Johannes M. Herrmann

## Abstract

Mitochondria are essential organelles that play a key role in cellular energy metabolism. Transitions between glycolytic and respiratory conditions induce considerable adaptations of the cellular proteome. These metabolism-dependent changes are particularly pronounced for the protein composition of mitochondria. Here we show that the yeast cytosolic ubiquitin conjugase Ubc8 plays a crucial role in the remodeling process when cells transition from respiratory to fermentative conditions. Ubc8 is a conserved and well-studied component of the catabolite control system that is known to regulate the stability of gluconeogenesis enzymes. Unexpectedly, we found that Ubc8 also promotes the assembly of the translocase of the outer membrane of mitochondria (TOM) and stabilizes its cytosol-exposed receptor subunit Tom22. Ubc8 deficiency results in a compromised protein import into mitochondria and a subsequent accumulation of mitochondrial precursor proteins in the cytosol. Our observations show that Ubc8, which is controlled by the prevailing metabolic conditions, promotes the switch from glucose synthesis to glucose usage in the cytosol and induces the biogenesis of the mitochondrial TOM machinery in order to improve mitochondrial protein import during phases of metabolic transition.

## Introduction

Mitochondria are of central relevance for cellular metabolism. They house the enzymes needed for respiration, the tricarboxylic cycle, the biogenesis of iron-sulfur clusters and multiple reactions in the synthesis and breakdown of amino acids, fatty acids, membrane lipids, ubiquinone and other metabolites (Wallace, 2005). Owing to their crucial role in metabolism, the volume and composition of the mitochondrial network is strongly responsive to changing metabolic conditions, much more so than that of most other cellular compartments (Morgenstern et al., 2017, Tsuboi et al., 2020). The shift from glycolytic fermentation to respiration in baker’s yeast is a well-studied metabolic transition which is accompanied by a major remodeling of the mitochondrial proteome (DeRisi et al., 1997, Di Bartolomeo et al., 2020). This metabolic transition is called the ‘diauxic shift’ and can be observed in glucose-grown yeast cultures. Cells initially produce ethanol by fermentative breakdown of glucose until glucose depletion induces a shift to respiration. A large fraction of the genes for mitochondrial proteins are induced by the diauxic shift, leading to a more than twofold increase in the total copy number of mitochondrial proteins (Morgenstern et al., 2017). A concerted transcriptional program leads to the induction of respiratory enzymes and other mitochondrial proteins. In parallel, the gene expression of gluconeogenic enzymes is induced to enable glucose replenishment by ethanol utilization (Galdieri et al., 2010, Laz et al., 1984, Liu and Barrientos, 2013, DeRisi et al., 1997). Thus, the ‘diauxic shift’ is mainly driven by responses on the transcriptional level.

The reverse transition from respiratory to fermenting conditions is far less understood and more complicated. In addition to changes in protein expression, many proteins, for example gluconeogenic enzymes or mitochondrial proteins, need to be reduced in their amount or even completely removed. This depletion is accomplished by a process termed catabolite degradation (Chiang and Schekman, 1991). The glucose-induced proteolysis of fructose-1,6-bisphosphatase (Fbp1) in ethanol-grown cultures was extensively studied and allowed the elucidation of the molecular mechanisms of catabolite degradation. A key factor in this remodeling process is the GID (Glucose Induced degradation Deficient) complex, which is conserved among eukaryotes and best studied in yeast. The components of this multi-subunit ubiquitin ligase (E3) were initially identified in a genetic screen for mutants deficient in glucose-induced Fbp1 degradation (Schüle et al., 2000, Hämmerle et al., 1998). A recently solved cryo electron microscopy structure of the GID complex showed that its 20 protein subunits resemble a large organometallic chelator with a central binding site for substrates and the ubiquitin conjugase (E2) Ubc8 (Sherpa et al., 2021). The conserved Ubc8 protein is homologous to other E2 enzymes (Qin et al., 1991) and seems to work exclusively in the context of GID-mediated protein degradation (Kong et al., 2021). Substrate binding to the constitutively expressed core of the GID complex occurs by substrate-specifying subunits whose expression depends on the prevailing metabolic conditions (Dong et al., 2018, Kong et al., 2021, Chen et al., 2017). Three substrate-specifying subunits have been identified: in the presence of glucose, Gid4 recruits gluconeogenesis enzymes such as Fbp1, phosphoenolpyruvate carboxykinase (Pck1), isocitrate lyase (Icl1) and cytosolic malate dehydrogenase (Mdh2). It thereby recognizes specific N-terminal proline motifs in their sequence (Chen et al., 2017). The substrate spectra and recognition motifs of the other specifying factors Gid10 (Melnykov et al., 2019) and Gid11 (Kong et al., 2021) are less well understood.

Here we show that Ubc8 has a second function in addition to its role in catabolite degradation. It promotes the biogenesis of the translocase of the outer membrane of mitochondria, the TOM complex, which serves as the general entry gate for mitochondrial precursor proteins (Araiso et al., 2019, Shiota et al., 2015). In the absence of Ubc8, the central outer membrane receptor Tom22 is diminished, and cells accumulate a partially assembled TOM complex of compromised function. Interestingly, previous studies already showed that the assembly of Tom22 is under metabolic control by different cytosolic kinases (Gerbeth et al., 2013). Our observations identify the E2 protein Ubc8 as an additional regulatory component in the biogenesis of the mitochondrial import machinery, demonstrating that the mitochondrial protein import system is under cooperative control of enzymes of the phosphorylation and the ubiquitination system.

## Results

### Ubc8-deficient mutants accumulate mitochondrial precursor proteins in the cytosol

Most mitochondrial proteins are synthesized in the cytosol as precursors and subsequently imported into mitochondria (Chacinska et al., 2009). Owing to the very short time between synthesis and import (Williams et al., 2014, Tsuboi et al., 2020), mitochondrial precursors only very transiently encounter the cytosol under physiological conditions. We recently developed a genetic screen to identify yeast mutants with slower import rates (Hansen et al., 2018). To this end, the orotidine-phosphate decarboxylase (Ura3) was expressed as a fusion protein with the mitochondrial protein Oxa1. Cell growth on uracil-deficient plates was only possible upon the cytosolic accumulation of the Oxa1-Ura3 fusion protein, which is promoted under conditions of impaired mitochondrial protein import (Fig. 1A, S1A). Using automated mating approaches, the Oxa1-Ura3 expression cassette was introduced into yeast libraries covering 4916 deletion mutants of nonessential genes as well as 1102 DAmP (decreased abundance by mRNA perturbation) mutants of essential genes (Schuldiner et al., 2005, Hansen et al., 2018). Eleven of these mutants showed robust growth on uracil-deficient plates and were described before (Hansen et al., 2018). Additionally, we observed several extra mutants with a more moderately increased uracil independence, including a strain lacking the ubiquitin conjugase Ubc8 (Fig. 1B, S1B, S1D). Loss of Ubc8 resulted in improved growth in uracil-deficient media indicating higher cytosolic levels of the Oxa1-Ura3 precursor (Fig. 1C, S1C).

**Fig. 1.**
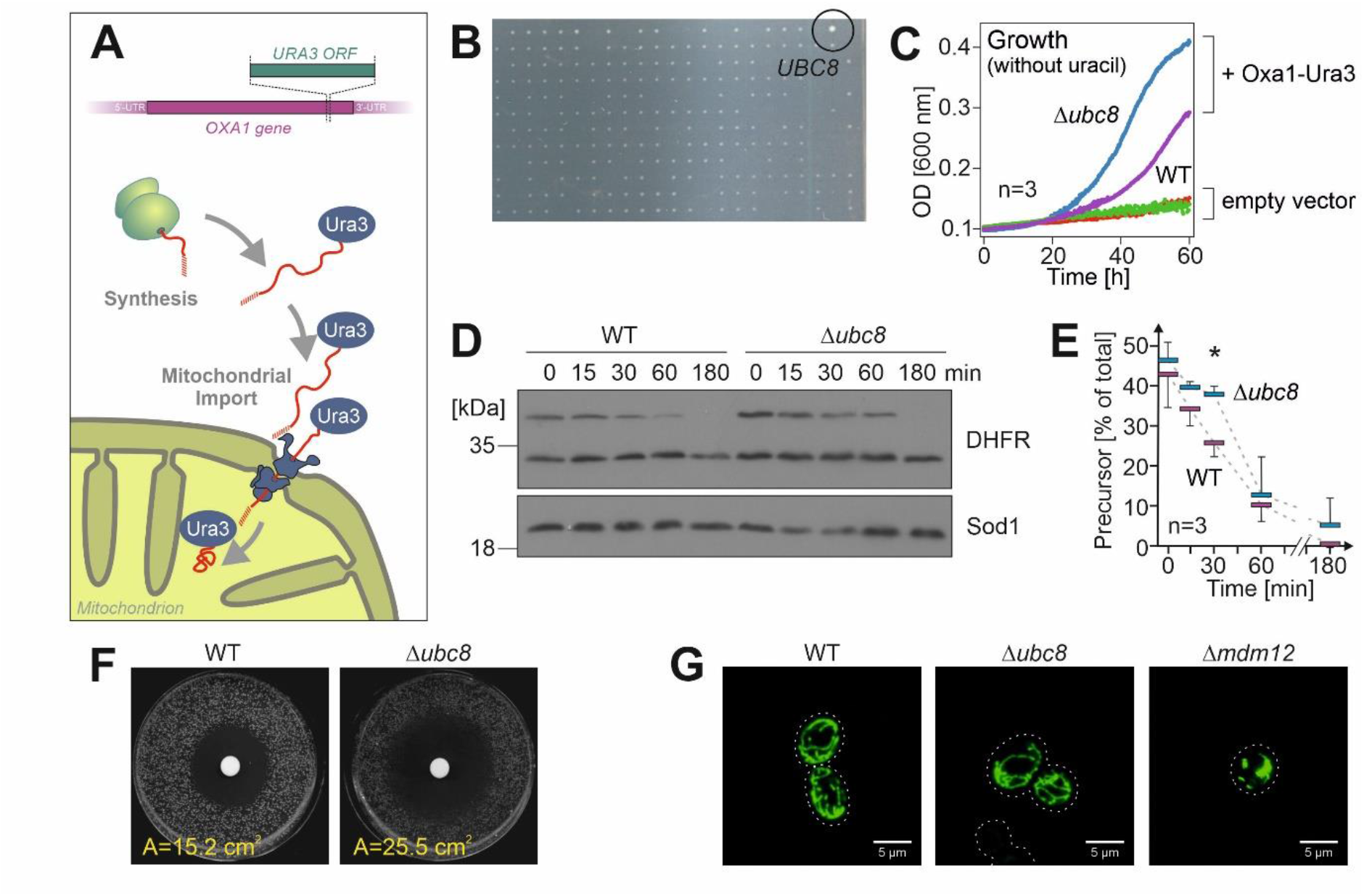
Loss of Ubc8 leads to the accumulation of mitochondrial precursor proteins in the cytosol. **A.** Schematic representation of the Oxa1-Ura3 screen. The accumulation of the Oxa1-Ura3 precursor in the cytosol confers uracil independence. **B.** Yeast deletion and DamP libraries with mutants expressing the Oxa1-Ura3 fusion protein were screened on uracil-deficient plates. Here a segment of a plate is shown after 1 d of growth at 30°C. Position of the Δ*ubc8* mutant is indicated. See Fig S1B for the entire plate. **C.** Wild-type (WT) and Δ*ubc8* cells harboring Oxa1-Ura3 expression plasmid or an empty vector for control were grown at 30°C for the times indicated in uracil-deficient synthetic media. Cell densities were measured. The graph shows mean values of three replicates. **D, E.** Wild type and Δ*ubc8* cells were transformed with a b_2_(1-167)_Δ19_-DHFR-expression plasmid. A galactose-grown preculture was shifted to galactose-free lactate medium for the times indicated. Aliquots were taken and analyzed by Western blotting using antibodies against DHFR and Sod1 for control. Panel E shows mean values and standard deviations of three biological replicates. **F.** Cells were spread on glycerol plates and a filter was placed to the center to which 10 µl 10 mM CCCP was added. Cells were incubated for one day and the inhibition area (A) was measured. **G.** A mitochondria-targeted GFP protein was expressed in the indicated cells (Dimmer et al., 2002). Cells were grown in galactose medium before the morphology of the mitochondrial structure was visualized by fluorescence microscopy. The Δ*mdm12* mutant was used as control for a strain with altered morphology (Dimmer et al., 2002).

Here, we asked whether the increased accumulation of the Oxa1-Ura3 precursor observed in Δ*ubc8* cells is indicative of a more general import problem that leads to the cytosolic accumulation of other mitochondrial precursors. To this end, we expressed the well characterized model protein b_2(1-167)Δ19_-DHFR in wild-type and *Δubc8* cells from a galactose-inducible *GAL* promoter. This fusion protein consists of a matrix targeting sequence followed by mouse dihydrofolate reductase (DHFR). The fast and rather stable folding of the DHFR domain results in a slow import and transient accumulation of the protein in the cytosol (Boos et al., 2019, Eilers and Schatz, 1986). When the galactose-driven expression of the fusion protein was stopped by a switch to galactose-free medium, the level of b_2(1-167)Δ19_-DHFR precursor declined over time. However, this decline occurred significantly slower in the *Δubc8* mutant than in wild-type cells (Fig. 1D, E). This indicates that in the absence of Ubc8, the precursor is either imported more slowly or degraded less efficiently than in wild-type cells. Consistent with these observations, *Δubc8* cells were hypersensitive to CCCP, which inhibits mitochondrial import by uncoupling the mitochondrial membrane potential (Fig. 1F). However, Ubc8 is not required for growth on respiratory media (Fig. S1E) (Qin et al., 1991) or the maintenance of mitochondrial morphology (Fig. 1G). In summary, our results indicate that the ubiquitin conjugase Ubc8 is directly or indirectly relevant for the depletion of mitochondrial precursor proteins from the cytosol, either by promoting their import or their proteasomal degradation (Fig. S1F).

### Ubc8 targets a divers set of substrates

We next employed mass spectrometry-based proteomics to gain a more comprehensive overview of the proteome dynamics in wild-type and Δ*ubc8* cells. The switch of media in a dynamic stable isotope labeling by amino acids in cell culture (SILAC) approach allows for a precise measurement of the turnover rates of individual proteins (Ong et al., 2002, de Godoy et al., 2008). This method proved to be very powerful to determine the import, assembly and degradation of mitochondrial proteins (Bogenhagen and Haley, 2020, Saladi et al., 2020, Schäfer et al., 2021). To this end, we grew wild-type and Δ*ubc8* cells in 2% lactate medium (which promotes respiration and gluconeogenesis) to mid-log phase with ‘light’ amino acids (i.e. [^14^N_2_, ^12^C_6_]-lysine, [^14^N_4_, ^12^C_6_]-arginine). After removing a first sample (t_0_), cells were harvested and either resuspended in ‘heavy’ (i.e. [^15^N_2_, ^13^C_6_]-lysine, [^15^N_4_, ^13^C_6_]-arginine) medium containing lactate or lactate plus 2% glucose (Fig. 2A). After growth for one doubling time, samples were taken and analyzed by mass spectrometry. Four independent replicates of each sample were analyzed, from which the data were processed and normalized as described in the Materials and Methods. Principle component analysis revealed that the proteome of the *Δubc8* and wild-type cells differ considerably even when cells are continuously grown in lactate, which suggests that Ubc8 is of relevance for respiring cells (Fig. 2B). Addition of glucose caused a further strong ‘catabolite effect’ on the proteomes of these cells.

**Fig. 2.**
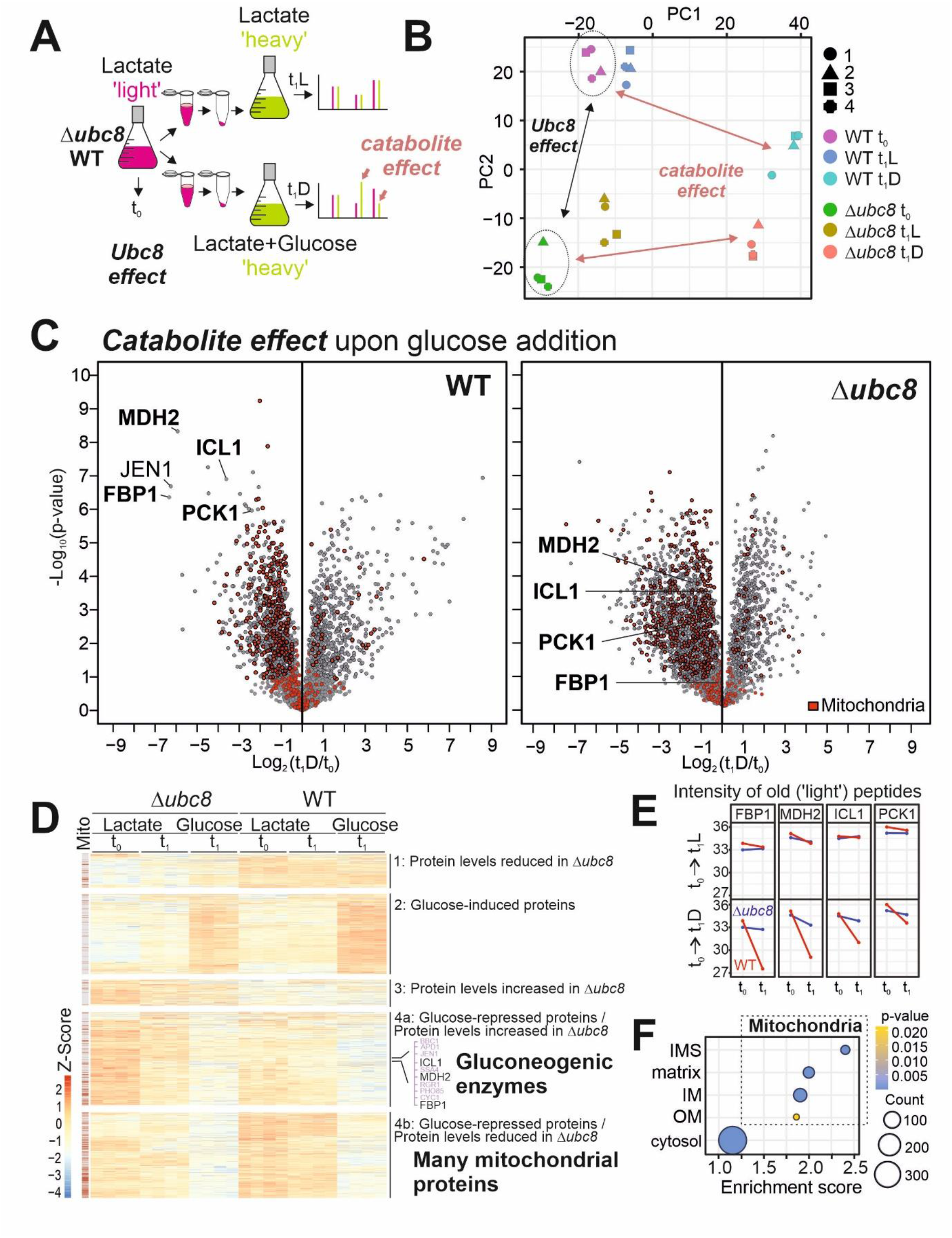
Ubc8 promotes catabolite degradation and influences levels of mitochondrial proteins. **A.** Schematics of the proteomics workflow. **B.** Principal component analysis of the data after normalization (see materials and methods). The entire dataset of the measurement is shown in Table S1. **C.** Volcano plot comparing the whole cell proteomes after and before the metabolic shift from lactate to glucose medium of WT and Δ*ubc8* cells. Positions of gluconeogenic enzymes regulated by Ubc8-dependent metabolic degradation are labeled in bold. Mitochondrial proteins are highlighted in red (Morgenstern et al., 2017). Corresponding plots for the samples that were further grown on lactate are shown as Fig. S2A. **D.** Hierarchical clustering of protein intensities identifies five distinct groups. Positions of mitochondrial proteins are indicated on the left. Gluconeogenic enzymes are found in group 4a. Group 4b contains many mitochondrial proteins. See Table S2 for additional information. **E.** Intensities in the light channel (‘old’ peptides) of the indicated proteins at t_0_ and t_1_ were plotted. Note, that lactate-to-glucose switches induces Ubc8-dependent degradation of gluconeogenic enzymes. **F.** Enrichment scores for proteins in group 4b indicate the presence of many components of mitochondrion-specific GO categories.

The glucose-induced shift from respiration to fermentation induces the depletion of gluconeogenic enzymes (Fbp1, Mdh2, Icl1 and Pck1) in wild-type but not in Δ*ubc8* cells (Fig. 2C), which is consistent with previous reports (Schüle et al., 2000, Hämmerle et al., 1998). In contrast, if cells were continuously grown in lactate, these enzymes did not differ considerably between wild-type and *Δubc8* cells (Fig. S2A).

Interestingly, we further noticed that the levels of many mitochondrial proteins were unexpectedly decreased in *Δubc8* cells. This was particularly obvious after hierarchical clustering of the proteomics data, which distinguished different groups of proteins according to differences in their abundance in lactate and glucose medium in wild-type and Δ*ubc8* cells, respectively (Fig. 2D, Suppl. Table S2). As expected, a defined cluster of gluconeogenesis enzymes showed a Ubc8-dependent decline upon glucose addition (Fig. 2D, group 4a, S2B). This was particularly apparent for the ‘light’ peptides, thus for old proteins, consistent with the regulation by proteolysis (Fig. 2E). In addition, a clustered group of other glucose-repressed proteins (Fig. 2D, group 4b) was defined by the fact that their levels were diminished in Δ*ubc8* cells even upon continuous growth in lactate (Fig. S2C). Interestingly, this group contained many mitochondrial proteins, indicating that the levels of many proteins of the IMS, the inner membrane and the matrix depend on the presence of Ubc8 (Fig. 2F, Supplemental Table S2). Thus, Ubc8 is not simply a glucose-induced removal factor of four gluconeogenesis enzymes but rather plays a much more general role in the adaptation of the cellular proteome to the prevailing metabolic state.

Ubc8 apparently has a second, distinct role as a factor that promotes the biogenesis of mitochondrial proteins. Interestingly, this latter function was not only apparent upon glucose supplementation to respiring cells but also observed when cells were continuously grown in lactate.

### Ubc8 is critical for metabolic remodeling of yeast cells

Ubc8 was initially discovered as a protein required for the rapid glucose-induced degradation of Fbp1 (Schüle et al., 2000). Consistent with these original reports, we observed that a shift from glycerol to glucose medium induces the rapid depletion of HA-tagged Fbp1 in an Ubc8-dependent manner (Fig. 3A). Such a Ubc8-mediated catabolite degradation was also observed for Icl1 and Mdh2 (Fig. 3B-D), thereby confirming previous studies (Karayel et al., 2020, Chen et al., 2021, Chen et al., 2017).

**Fig. 3.**
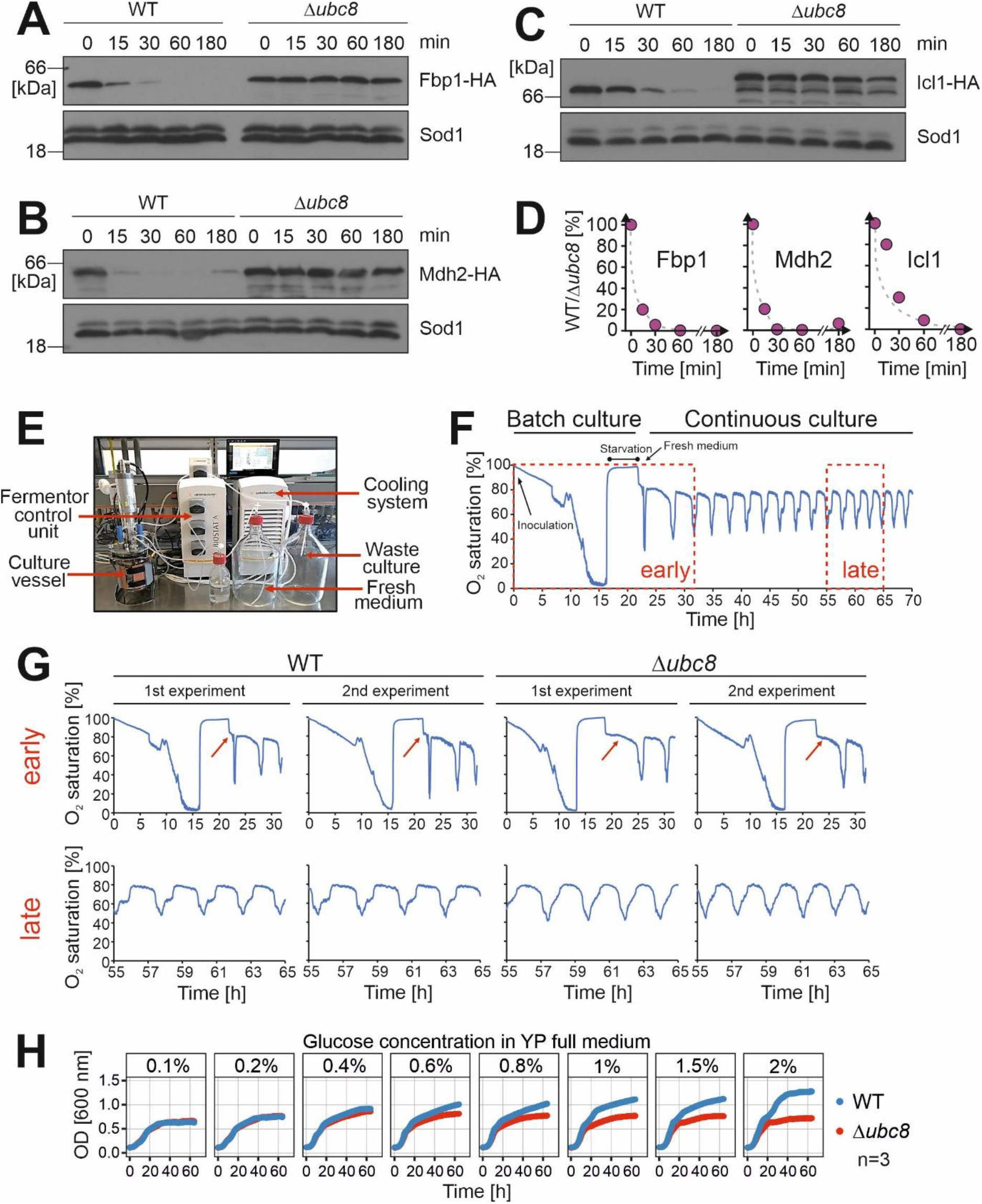
Ubc8 is crucial for the quick adaptation to metabolic changes. **A-D.** Wild-type and Δ*ubc8* cells expressing the indicated HA-tagged proteins under endogenous control were cultured in lactate medium. The medium was replaced by glucose medium. Aliquots were taken after time points indicated and analyzed by Western blotting. Sod1 served as loading control. Panel D shows the quantification of A-C. **E**. Setup of the fermentor system. **F**, **G**. The fermentor run was initiated by inoculation of a YPD-grown preculture. Oxygen saturation in the medium was automatically monitored every 10 s. Cells grew as a batch culture until stationary phase. The culture was then kept in starvation phase for 5 h to synchronize all cells. Finally, fresh glucose-medium was continuously added at defined rates to initiate metabolic cycling of the cultures. Early and late phases of the runs are shown for wild-type and Δ*ubc8* cultures. **H**. Cultures were grown to mid-log phase in glucose medium and diluted in medium containing different concentrations of glucose as indicated. Cell densities were continuously measured during three days of growth under continuous shaking.

Even though the role of Ubc8 in catabolite degradation of Fbp1 and other gluconeogenic enzymes is well documented, little is known about the physiological relevance of Ubc8 (and the GID complex). Since yeast cells are typically cultured under static growth conditions, the adaptation to varying metabolic conditions is normally of little relevance. However, under defined growth conditions in a continuous fermentor-based culture yeast cells oscillate between phases of low and high oxygen consumption in a phenomenon termed the ‘yeast metabolic cycle (YMC)’ (Slavov et al., 2011, Tu et al., 2005, Amponsah et al., 2021). After initiation of a fermentor run by injection of a glucose-grown preculture, yeast cells deplete glucose and oxygen before entering a starvation phase (Fig. 3E, F). When fresh glucose-containing medium is pumped into the vessel, a spontaneous and stable oscillation of the oxygen saturation in the culture medium is induced. Importantly, oxygen saturation in the medium is a direct consequence of a population-synchronized oscillation in oxygen consumption (Tu et al., 2005). Periodic changes in oxygen consumption in continuous and synchronized yeast cultures are the most obvious and distinctive feature of the YMC. To test for the relevance of Ubc8 in this context, we deleted the *UBC8* gene in CEN.PK 113-1A yeast strain (Amponsah et al., 2021, Burnetti et al., 2016) and monitored oxygen consumption in the fermentor for several days. Thereby, we noticed that whilst both wild-type and Δ*ubc8* cultures exhibited the characteristic oxygen saturation cycling, the cycles in the Δ*ubc8* strain only started after a four hour delay, that was not observed in wild-type cells (Fig. 3G, read arrows). Furthermore, the cycles in Δ*ubc8* cells tend to be shorter (Fig. 3G, lower panels), which is consistent with a disrupted ability to switch between different metabolic states. Still, Δ*ubc8* cells are able to alternate rhythmically between oxygen-consuming and fermenting phases.

Next, we tested the growth of cells after switching from glycerol medium to different concentrations of glucose (Fig. 3H). We observed no difference in the growth of wild-type and Δ*ubc8* cells when the glucose concentrations were low. This was surprising because we expected that the presence of futile cycles, owing to the simultaneous presence and activity of glycolytic and gluconeogenic enzymes in Δ*ubc8* cells, would waste energy and should negatively affect cell growth, particular when carbon sources are scarce. However, at higher glucose concentrations, wild-type cells grew to much higher cell densities than Δ*ubc8* cells did which was not caused by the production of toxic byproducts such as methylglyoxal (Fig. S3A). We noticed that the growth curves of wild-type and Δ*ubc8* cells were identical for the first few hours but the Δ*ubc8* cells lagged behind after the diauxic shift, i.e. the switch from fermentative ethanol production to ethanol-consuming respiration. The diauxic shift is accompanied by dramatic rearrangements of mitochondrial function and structure in yeast cells (Di Bartolomeo et al., 2020). Furthermore, we noticed increased lethality of Δ*ubc8* cells in stationary, glucose-grown cultures (Fig. S3B), consistent with problems under metabolic conditions that rely on mitochondrial activity (Ocampo et al., 2012). Thus, these physiological data indicate that the deletion of *UBC8* shows consequences that point to problems in mitochondrial functionality.

### Efficient TOM assembly depends on Ubc8

Due to the reduced functionality of mitochondria in Δ*ubc8* cells and the increased amounts of mitochondrial precursor proteins in the cytosol, we directly tested whether Ubc8 is relevant for protein import into mitochondria. To this end, we isolated mitochondria from wild-type and Δ*ubc8* cells and performed *in vitro* import reactions with the inner membrane protein Oxa1 (which uses the TOM-TIM23 import pathway) and the ATP/ADP carrier Pet9 (which embarks on the TOM-TIM22 import route). Both proteins were imported with considerably reduced efficiency into Δ*ubc8* mitochondria (Fig. 4A), indicating that Ubc8 is important for the biogenesis or stability of the mitochondrial protein import system.

**Fig. 4.**
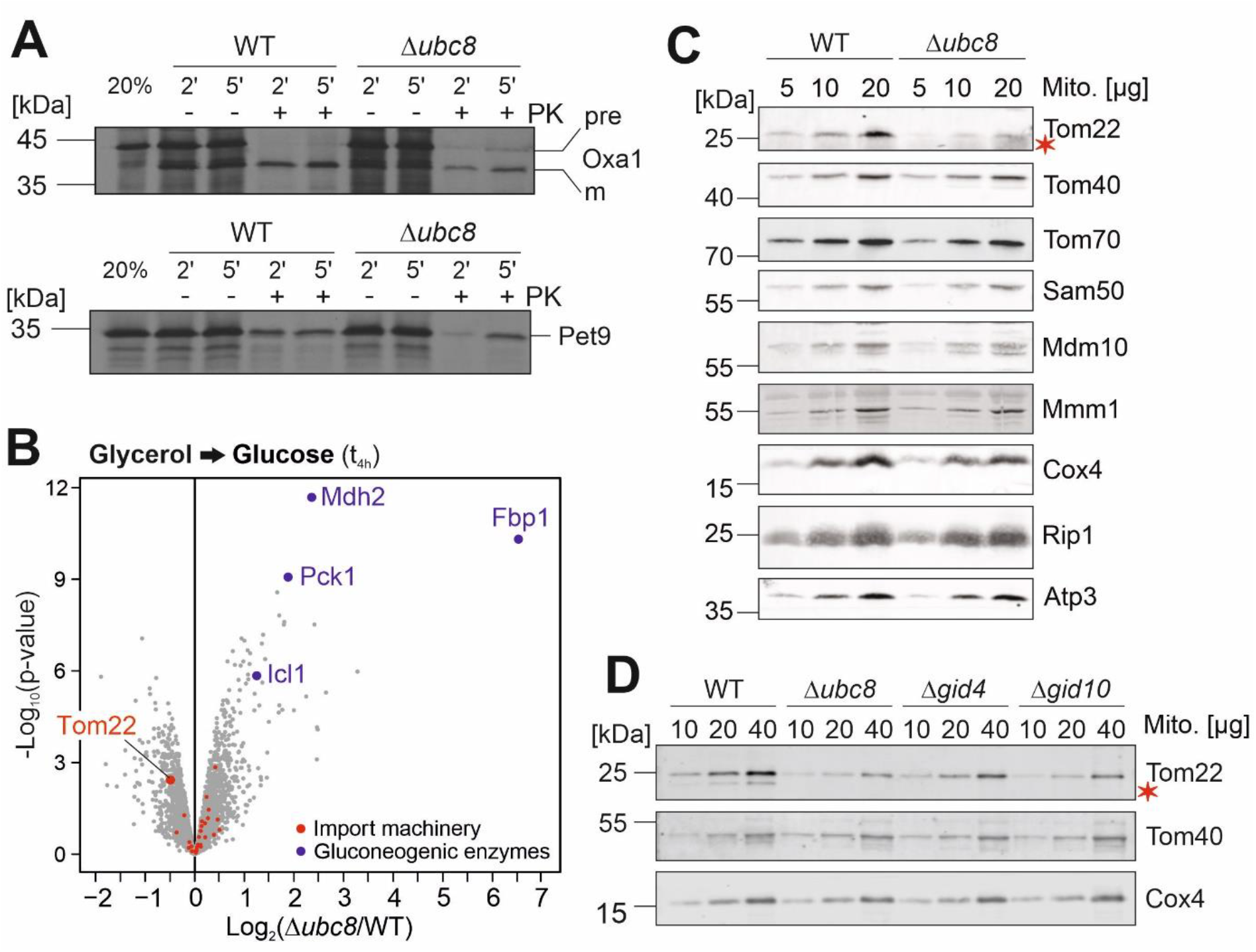
Absence of Ubc8 or other components of the GID complex leads to diminished levels of the outer membrane protein Tom22. **A.** Radiolabeled Oxa1 and Pet9 were incubated with mitochondria isolated from wild-type or Δ*ubc8* cells at 30°C for different times. Non-imported proteins were degraded by addition of proteinase K (PK). 20% of the radiolabeled proteins used per import reaction are loaded for control. Proteins were visualized by autoradiography. **B.** Cells were grown in glycerol medium before glucose was added for 4 h. Cells were subjected to mass spectrometry and the levels of different groups of mitochondrial proteins were analyzed. Four replicates were analyzed. See also Fig. S4A, B. and Table S3. **C, D.** Cells were grown in glycerol medium. After addition of glucose for 4 h, mitochondria were isolated and subjected to SDS PAGE. The indicated proteins were visualized by Western blotting.

To identify the reason for the import defect of Δ*ubc8* cells, we shifted wild-type and Δ*ubc8* cells from glycerol to glucose medium for 4 h and analyzed their proteomes by mass spectrometry (Fig. 4B, S4A, B. See also Supplemental Table S3). After the metabolic shift, but not before, Δ*ubc8* cells showed considerably increased levels of gluconeogenic enzymes, as expected. In addition, we noticed that the absence of Ubc8 caused diminished levels of the mitochondrial protein Tom22, which was less pronounced in glycerol-grown cells.

Western blot experiments of isolated mitochondria confirmed the reduced Tom22 levels (Fig. 4C), whereas the levels of the pore-forming subunit of the TOM complex, Tom40, and also that of the receptor Tom70, remained unaffected. Mitochondria isolated from Δ*gid4* and Δ*gid10* cells also showed reduced Tom22 levels. Gid4 and Gid10 serve as substrate-binding subunits of the GID complex (Chen et al., 2017, Melnykov et al., 2019, Sherpa et al., 2021). Thus, the GID complex, for which Ubc8 serves as the ubiquitin conjugase, increases the levels of the crucial TOM protein Tom22 in mitochondria, which is a rate-limiting factor of the mitochondrial protein import system (Schmidt et al., 2011, Zeng et al., 2019, Shiota et al., 2011, van Wilpe et al., 1999).

### Ubc8 facilitates the assembly of Tom22 into functional TOM complexes

Next, we analyzed the size of the TOM complex by blue native gel electrophoresis followed by Western blotting (Fig. 5A). In wild-type mitochondria, the TOM complex migrates like the 440 kDa molecular weight marker. In contrast, in Δ*ubc8* mitochondria, Tom40 and Tom22 were part of a faster migrating complex. Again, we observed considerably reduced levels of Tom22. The sizes of other complexes containing other mitochondrial proteins (Sam50 or Cox1) were not altered.

**Fig. 5.**
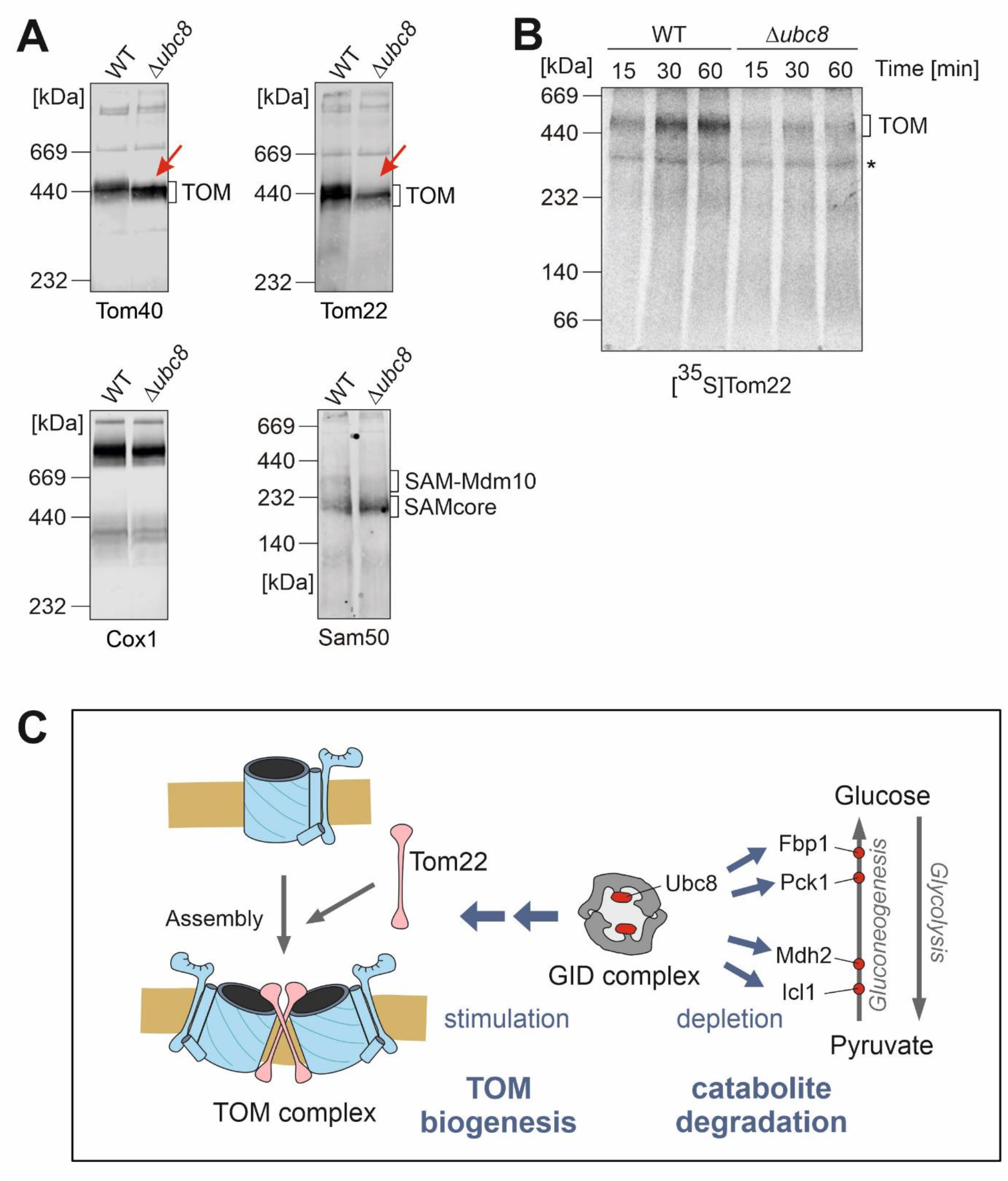
Ubc8 is required for full assembly of the TOM complex. **A.** Cells were grown in glycerol medium. After addition of glucose for 4 h, mitochondria were isolated and subjected to blue native gel electrophoresis. The indicated proteins were visualized by Western blotting. **B.** Radiolabeled Tom22 was incubated with mitochondria isolated from wild-type and Δ*ubc8* cells for the time indicated. Assembly of Tom22 into the TOM complex was analyzed by blue native gel electrophoresis and autoradiography. The position of the TOM complex is indicated. Note that the assembly of Tom22 into Δ*ubc8* mitochondria occurs with reduced efficiency. **C.** Model of Ubc8 function. See text for details.

The assembly of Tom22 with other subunits of the TOM complex is well characterized. The efficiency of Tom22 assembly depends on a complex regulatory network of cytosolic kinases and phosphatases (Rao et al., 2012, Schmidt et al., 2011, Gerbeth et al., 2013, Kravic et al., 2018) and its interaction with several integral factors of the mitochondrial outer membrane (Sakaue et al., 2019, Ellenrieder et al., 2016, Wu et al., 2018, Becker et al., 2011, Qiu et al., 2013, Meisinger et al., 2004, Keil and Pfanner, 1993). In order to test which of these steps are Ubc8-dependent, we imported radiolabeled Tom22 into mitochondria isolated from wild-type and Δ*ubc8* cells and monitored its assembly into the TOM complex by blue native gel electrophoresis (Fig. 5B). Thereby, we noticed that Tom22 assembly was decreased in the absence of Ubc8, indicating that the reduced Tom22 levels were due to impaired biogenesis of Tom22.

In conclusion, the presence of Ubc8 is obviously important to facilitate the biogenesis of the TOM complex, particularly after metabolism switches from respiration to fermentation conditions. If Ubc8 is absent, only a partially assembled and functionally compromised TOM complex is formed, which leads to a moderate, but wide-ranging depletion of mitochondrial proteins, in particularly those of the matrix for which the Tom22 levels seem particularly important (Fig. S4C). Thus, biogenesis of the major protein entry site of the mitochondria, the TOM complex, is under control of the ubiquitin conjugase Ubc8 (Fig. 5C). To our knowledge this is the first observation that the cytosolic ubiquitin system controls the biogenesis of the mitochondrial import machinery.

## Discussion

Cells dynamically shape their proteome in response to the prevailing growth conditions. In particular, metabolic changes can induce a massive reorganization of the cellular proteome (Nielsen, 2017). For example, in yeast cells, the addition of glucose to galactose-grown cells changes the expression of 25% of all gene products by more than twofold, even though neither growth rates nor cell sizes are affected (Ronen and Botstein, 2006). In a nutshell, proteome remodeling is the result of altered gene expression on the one hand and regulated protein degradation on the other. Thus, the orchestrated interplay of transcription factors and components of the ubiquitin-proteasome system (UPS) is crucial for dynamically and accurately adapting the proteome.

We designed a quantitative proteomics experiment based on dynamic SILAC labeling (Pratt et al., 2002) to follow the changes on the yeast proteome induced by switches from respiratory glucose synthesis to fermentative glucose consumption. Dynamic SILAC labeling has been successfully used in the past to measure the rates of protein turnover on the basis of the relative loss of ‘old’ (in our case light) peptides (Mathieson et al., 2018, Doherty et al., 2009, Dorrbaum et al., 2018, Saladi et al., 2020) or the rates of protein synthesis based on the accumulation of ‘new’ (in our case heavy) peptides (Schäfer et al., 2021). Our results document the potential of metabolic labeling for both aspects:

(1) On the one hand, we observed a rapid Ubc8-dependent depletion of the gluconeogenic enzymes upon glucose addition, resulting in the rapid loss of ‘old’ peptides of Fbp1, Mdh2 and Icl1in wild-type but not in Δ*ubc8* cells. This is consistent with the established role of Ubc8 in catabolic degradation of gluconeogenic enzymes (Schüle et al., 2000, Chen et al., 2017, Chen et al., 2021, Karayel et al., 2020, Sherpa et al., 2021). Even though Δ*ubc8* cells had considerably higher levels of gluconeogenic enzymes, their growth rates remained unaffected even when glucose levels were low. Thus, futile cycles, caused by the simultaneous presence of glycolytic and gluconeogenic enzymes, might either be also prevented by other means such as posttranslational modifications and allosteric regulation of the enzymes (Smets et al., 2010, Dihazi et al., 2003), or do not directly impair growth.

(2) On the other hand, we unexpectedly detected that the absence of Ubc8 strongly impairs the biosynthesis of mitochondrial proteins, proposing a role of Ubc8 in mitochondrial biogenesis. The dynamic SILAC method made it possible to discriminate between changes in protein expression and those in protein stability within the same experiment. In this study, we confirmed the role of Ubc8 for the general accumulation of mitochondrial proteins by a second proteomics experiment using a simpler label-free quantification procedure (see Fig. 4B) in which the quantification was more straight-forward. However, this second experiment did not allow us to tell whether mitochondrial proteins were less produced or faster degraded.

The absence of Ubc8 considerably reduced the levels of the mitochondrial outer membrane protein Tom22. Previous studies by Chris Meisinger and coworkers described a glucose-dependent regulation of the import and assembly of Tom22 in depth (Schmidt et al., 2011, Gerbeth et al., 2013, Kravic et al., 2018). In the cytosol, Tom22 is phosphorylated by the casein kinases CK1 and CK2 as well as by the protein kinase A (PKA). Whereas CK1/2-mediated phosphorylation promotes Tom22 import, phosphorylation by PKA was found to reduce the association of Tom22 with the TOM complex. The phosphatases counteracting Tom22 phosphorylation were not identified so far. It is conceivable that Ubc8 mediates the degradation of the phosphatase that counteracts CK1/2-mediated degradation or, alternatively, the degradation of PKA. In both cases, reduced Tom22 levels are expected. Indeed, in human cells the GID complex serves as the ubiquitin ligase of PKA (Liu et al., 2020).

Apparently, Tom22 represents the critical key element in the regulation of mitochondrial protein import. What makes Tom22 so special? Tom22 is a central component of the TOM complex that serves as ‘multifunctional organizer’ (van Wilpe et al., 1999). This term appropriately describes the function of Tom22, as in the absence of Tom22, the TOM complex dissociates in core units that mainly consist of the beta barrel protein Tom40. The addition of Tom22 converts these pores into fully functional dimeric and trimeric TOM complexes thereby considerably enhancing protein translocation efficiency (van Wilpe et al., 1999, Araiso et al., 2019, Shiota et al., 2015, Zeng et al., 2019, Tucker and Park, 2019, Bausewein et al., 2017, Wang et al., 2020). Recent studies indicated that Tom22 association drives a dynamic conversion of different functional states of the TOM complex, which modulates its substrate preference (Sakaue et al., 2019, Gornicka et al., 2014). Thus, the controlled incorporation of Tom22 into the TOM complex seems to be an excellent strategy to modulate mitochondrial import capacity on a general scale (Araiso et al., 2021). Interestingly, while Ubc8 induces Fbp1 degradation exclusively upon glucose addition, Ubc8 improved Tom22 assembly also under continuous non-fermentative conditions. This suggests that the Ubc8 effect on mitochondrial protein import is not exclusively mediated via the glucose-controlled substrate-specifying factor Gid4. It appears likely that Ubc8 and the GID complex control the assembly of the TOM complex also via other substrate-specifying factors, potentially via Gid10. It will be exciting in the future to unravel the molecular details by which the phosphorylation and the ubiquitination systems in the cytosol cooperate to control mitochondrial protein biogenesis.

## Materials and Methods

### Yeast strains and plasmids

All yeast strains used in this study are based on the wild-type strains BY4742 (MATα *his3 leu2 lys2 ura3*), YPH499 (MATa *ura3 lys2 ade2 trp1 his3 leu2*), YPH499 *Δarg4* (MATa *ura3 lys2 ade2 trp1 his3 leu2 arg4*) or CEN.PK113.1A (MATα)(Sikorski and Hieter, 1989). To delete *UBC8* and *GID10*, the genes were chromosomally replaced by kanMX4 or natNT2 cassettes using pFA6α-kanMX4 and pFA6α-natNT2 as templates, respectively. For HA-tagging of Fbp1, Mdh2, and Icl1, the sequence of 6HA-natNT2 was amplified from the plasmid pYM17 and genomically integrated downstream of the respective genes (Janke et al., 2004). BY4742 deletion strain for *GID4* was taken from a yeast deletion library (Giaever et al., 2002). Yeast strains used in this study are listed in the Supplemental Table S5.

For yeast transformation the lithium acetate method was used (Gietz et al., 1992). Oxa1-Ura3 and b_2_Δ_19(1-167)_-DHFR plasmids were described previously (Hansen et al., 2018, Boos et al., 2019). Empty plasmids were used for control.

Yeast cells were grown in yeast full medium containing 1% (w/v) yeast extract, 2% (w/v) peptone and 2% of the respective carbon source at 30°C. As carbon sources glucose (YPD), galactose (YPGal) or glycerol (YPG) were used, as indicated. Strains carrying plasmids were grown in minimal synthetic medium containing 0.67% (w/v) yeast nitrogen base and 2% of glucose (SD), galactose (SGal) or lactate (SLac), as indicated. For plates, 2% of agar was added to the medium. To induce protein expression from the *GAL1* promotor, 0.5% of galactose was added to the yeast cultures.

### Growth assays and viability test

For drop dilution assays, respective yeast strains were grown in yeast full or synthetic media to mid-log phase. After harvesting 0.5 OD_600_ of cells and washing with sterile water, a 1:10 dilution series was prepared. 3 µl of each dilution were dropped on agar plates containing full or synthetic media. Pictures of the plates were taken after different days of incubation at 30°C. For testing growth in liquid media, growth curves were performed in 96-well plates in technical triplicates. The ELx808 Absorbance Microplate Reader (BioTek) was used for automated OD measurement at 600 nm. With starting at an OD_600_ of 0.1, OD_600_ was measured every 10 min for 72 h at 30°C.

To test the viability of cells, respective yeast strains were inoculated in YPD medium, diluted once and then continuously grown in YPD for 10 days at 30°C. Samples were taken every day. For this, 1 OD_600_ of cells was harvested, washed with sterile water and 100 µl of OD_600_ 0.01 were plated out on YPD agar plates. After incubating for one to two days at 30°C, colony numbers were counted.

### Halo assay for CCCP sensitivity

For the halo assay, yeast strains were precultured in YPG medium. After harvesting 1 OD_600_ of cells and washing with sterile water, 100 µl of OD_600_ 0.01 were plated out on glycerol plates. A filter plate with 10 µl of CCCP (10 mM) was placed in the middle of the plate. After incubation for one day at 30°C, halo areas were measured. Filter plates with 10 µl DMSO served as negative controls.

### Preparation of whole cell lysates

4 OD_600_ of yeast cells were harvested and washed with sterile water. Pellets were resuspended in 40 µl/OD_600_ Laemmli buffer containing 50 mM DTT. Cells were lysed using a FastPrep-24 5 G homogenizer (MP Biomedicals) with 3 cycles of 20 s, speed 6.0 m/s, 120 s breaks, glass beads (Ø 0.5 mm) at 4°C. Lysates were heated at 96°C for 5 min and stored at −20°C until visualization by Western blotting.

### Antibodies

Antibodies were raised in rabbits using recombinant purified proteins. The secondary antibody was obtained from Bio-Rad (Goat Anti-Rabbit IgG (H+L)-HRP Conjugate #172-1019). The horseradish-peroxidase coupled HA antibody was purchased from Roche (Anti-HA-Peroxidase, High Affinity (3F10), #12 013 819 001). Antibodies were diluted in 5% (w/v) nonfat dry milk in 1x TBS buffer with the following dilutions: Anti-HA 1:500 and Anti-Rabbit 1:10,000. Antibodies used in this study are listed in Supplemental Table S4.

### Fermentor measurements

To measure population-synchronized metabolic oscillations, yeast cultures were stably synchronized with respect to the yeast metabolic cycle as previously described (Tu et al., 2005, Amponsah et al., 2021). All experiments were performed in a CEN.PK113-1A strain background. A Biostat A fermentor (Sartorius Stedim Systems), with a culture volume of 800 ml was used for all experiments. Culture media consisted of 10 g/l glucose, 1 g/l yeast extract, 5 g/l (NH_4_)_2_SO_4_, 2 g/l KH_2_PO_4_, 0.5 g/l MgSO_4_, 0.1 g/l CaCl_2_, 0.02 g/l FeSO_4_, 0.01 g/l ZnSO_4_, 0.005 g/l CuSO_4_, 0.001 g/l MnCl_2_, 2.5 ml 70% H_2_SO_4_ and 0.5 ml/l Antifoam 204.

Fermentor runs were initiated by the addition of a 20 ml preculture, which was grown to stationary phase in YPD medium at 30°C. The fermentor was run in batch-culture mode at 30°C, with an aeration rate of 1 l/min and constant stirring at 530 rpm. A constant pH of 3.4 was maintained by the automated addition of 10% (w/v) NaOH. Fermentor cultures were grown until ∼5 h after the exhaustion of carbon source as determined by continuous monitoring of culture oxygen saturation. Subsequently, a continuous culture was initiated by the addition of fresh media to the culture vessel at a dilution rate of 0.05 h^-1^. Culture oxygen saturation was automatically recorded with a sampling interval of 10 s.

### Sample preparation and mass spectrometric identification of proteins

For dynamic SILAC mass spectrometry (MS), YPH499 *Δarg4* wild-type and Δ*ubc8* cells were cultured in SLac medium containing light [^14^N_2_, ^12^C_6_]-lysine and [^14^N_4_, ^12^C_6_]-arginine isotopes. Cells were diluted continuously to keep them in the exponential growth phase while increasing the culture volume stepwise up to 300 ml. For timepoint t(0) 100 ml of each culture were harvested (5,000 *g*, 5 min, RT), washed with sterile water, and shock frozen with liquid nitrogen. Samples were stored at −80°C for further analysis. To analyze the metabolic shift (Fig.2, S2), 2×100ml of every culture was harvested (5,000 *g*, 5 min, RT) and washed with 30 ml SLac medium without lysine and arginine (5,000 *g*, 5 min, RT). Cells were resuspended in 200 ml SLac+2% glucose or SLac as control. Both media only contained the heavy amino acid isotopes of lysine (^15^N_2_, ^13^C_6_) and arginine (^15^N_4_, ^13^C_6_). Cultures were incubated at 30°C and 140 rpm for one doubling time of the cells. Cell growth was continuously monitored by measuring OD_600_. Samples (t(1) Gluc and t(1) Lac) were harvested and treated like described before for the samples t(0).

MS samples were prepared according to a published protocol with minor adaptations (Kulak et al., 2014). Cell lysates from 25 OD were prepared in 100 µl lysis buffer (6 M guanidinium chloride, 10 mM TCEP-HCl, 40 mM chloroacetamide, 100 mM Tris pH 8.5) using a FastPrep-24 5 G homogenizer (MP Biomedicals) with 3 cycles of 20 s, speed 8.0 m/s, 120 s breaks, glass beads (Ø 0.5 mm) at 4°C. Samples were heated for 10 min at 96°C and afterwards centrifuged twice for 5 min at 16,000 g. In between the supernatant was transferred to fresh Eppendorf tubes to remove all remaining glass beads. Protein concentrations were measured using the Pierce BCA Protein Assay (Thermo Scientific, #23225). For protein digestion, 36 µg of protein was diluted 1:10 with LT-digestion buffer (10% acetonitrile, 25 mM Tris pH 8.8). Trypsin (Sigma-Aldrich #T6567) and Lys-C (Wako #125-05061) were added to the samples (1:50 w/w). Samples were incubated over night at 37°C and 700 rpm. After 16 h, fresh Trypsin (1:100 w/w) was added for 30 min (37°C, 700 rpm). pH of samples was adjusted to pH <2 with trifluoroacetic acid (10%) and samples were centrifuged for 3 min at 16,000 g and RT. Desalting/mixed-Phase cleanup with 3 layer SDB-RPS stage tips (cat 2241). Samples were dried down in speed-vac and resolubilized in 12 µl buffer A++ (buffer A (0.1% formic acid) and buffer A* (2% acetonitrile and 0.1% trifluoracetic acid) in a ratio of 9:1)).

To analyze the metabolic shift by label-free mass spectrometry (Fig. 4B, S4B) the respective yeast strains were cultured in YPG. Glucose (final concentration: 2%) was added to the cultures for 4 h. 10 OD_600_ of cells were harvested (5,000 *g*, 5 min, RT) and washed with sterile water. Pellets were snap frozen in liquid nitrogen and stored at −80°C for further analysis. Samples were prepared for mass spectrometry as described above with minor changes: Cell lysates of 10 OD were prepared in 200 µl lysis buffer and 25 µg of protein was used for trypsin and Lys.C digestion.

For both MS experiments, digested peptides were separated on reversed-phase columns (50 cm, 75 μm inner diameter packed in-house with C18 resin ReproSilPur 120, 1.9 μm diameter (Dr. Maisch) using an Easy-nLC 1200 system (Thermo Scientific) directly coupled to a Q Exactive HF mass spectrometer (Thermo Scientific). A 3 hour gradient from 5% - 95% Solvent B (Solvent A: aqueous 0.1% formic acid; Solvent B: 80 % acetonitrile, 0.1% formic acid) at a constant flow rate of 250 nl/min was used to elute bound peptides. Further details of the gradient and instrument parameters are provided with the raw files uploaded to the ProteomeXchange repository.

MS data were processed using the MaxQuant software (version 1.6.10.43) (Cox and Mann, 2008, Cox et al., 2011, Tyanova et al., 2016) and a *Saccharomyces cerevisiae* proteome database obtained from Uniprot.

### Quantification and statistical analysis

MaxQuant output files were processed using Perseus and the R programming language. Each condition was measured in four replicates. For the lactate to glucose shift experiment (Fig 2), proteins were filtered to contain at least 3 valid values in at least one of compared conditions. Log2 protein intensities (combined intensity of the heavy and light channel) were mean centered. For the dynamic SILAC experiment (Fig. 2E), the same normalization factors were applied to the individual light and heavy channel. To compare conditions, a Student’s *t*-Test was performed and p-values were adjusted for multiple testing (Benjamini and Hochberg, 1995). Hierarchical clustering (Fig. 2D) was carried out in Perseus. Mean centered log2 protein intensities were filtered using a multiple-sample ANOVA test (FDR > 0.01, S0=1) implemented in Perseus and Z-scored protein intensities were clustered according to Euclidean distance. The heatmap was visualized in R using the pheatmap package. Gene set enrichment analysis was performed using Fisher’s exact test. *p-*values of gene set enrichments were adjusted for multiple hypothesis testing using the Benjamini-Hochberg procedure (Benjamini and Hochberg, 1995).

For the label-free experiment (Fig. 4 and S4) proteins were filtered as described above and label-free quantitation (LFQ) protein intensities were cleaned for batch effects using limma (Ritchie et al., 2015) and further normalized using variance stabilization normalization (Huber et al., 2002). Proteins were tested for differential expression using the limma package and *p-*values were adjusted for multiple hypothesis testing using the Benjamini-Hochberg procedure (Benjamini and Hochberg, 1995).

Western blot analyses were independently replicated with similar results, and representative data are shown in the figures. Quantification was performed with Fiji/ImageJ and significance testing was performed with Student’s *t*-Test.

### Sample preparation and mass spectrometric measurement of methylglyoxal (MG) levels in yeast

Respective yeast strains were cultured in yeast full medium containing 0.2% glucose until mid-log phase at 30°C. Glucose (final concentration 2%) was added to cultures for 1 h. For mass spectrometric analysis, 50 OD_600_ of cells were harvested (5,000 *g*, 5 min, RT). Cell pellets were washed with sterile water twice (5,000 *g*, 5 min, RT and 16,000 *g*, 2 min, RT respectively) and stored at −80°C for further analysis. MG levels were determined by mass spectrometry after derivatization to 2-methylquinoxaline (2-MQ). For sample derivatization, preparation of standards and mass spectrometric measurement, the protocol from Rabbani and Thornalley was adjusted for yeast cells (Rabbani and Thornalley, 2014). In detail: cells were resuspended in 300 µl ice-cold trichloroacetic acid (TCA, 20% (w/v) with 0.9% (w/v) sodium chloride). Glass bead lysis was performed using a FastPrep-24^TM^ 5G (MP Biomedicals) with 3 cycles of 20 s, speed 8.0 m/s, 120 s breaks, glass beads (Ø 0.5 mm) at 4°C. Samples were centrifuged (10,000 *g*, 10 min, 4 °C) and 140 µl of supernatant was used for derivatization. Sample derivatization was performed as described by Rabbani and Thronalley but quadruplicate volume of chemicals and d_4_-2-methylquinoxaline (d_4_-2-MQ) instead of isotopically labeled MG was used. Nine standards with different amounts of MG were prepared for a calibration curve (0-1091 nM).

For analysis, an Agilent 1200 HPLC system (Agilent Technologies, Waldkirch, Germany) coupled with an API 3200 tandem mass spectrometer (AB Sciex, Darmstadt, Germany) was used. Separations were performed using a RP 18 column (Xbridge, C18, 2.5 µm, 2.1 x 50 mm, Waters, Milford, USA). The injection volume was 15 µL and the used flow was 250 µL/min. The mobile phases consisted of water with 0.1% trifluoroacetic acid (TFA) (A) and a mixture of water with acetonitrile (50/50 (v/v)) with 0.1% TFA (B). Concentration of B was 0% at 0 min and increased to 100% for 5 min and is held at 100% for 5 min, followed by a reconditioning step. The measurement was carried out in the multiple reaction monitoring (MRM) mode with positive ionization. An electrospray ionization (ESI) source was used with source parameters as follows: ion spray voltage 5000 V, temperature 650°C, nebulizer gas 65 psi, heater gas 65 psi. The characteristic combinations of the parent ions and the product ions (Q1→Q3, *m/z*) for 2-MQ and d_4_-2-MQ were 145.1→77.1*, 145.1→92.1 and 149.1→81.1*, 149.1→122.1, respectively. For quantification a calibration curve with peak area ratio of 2-MQ/d_4_-2-MQ, where the transitions marked with an asterisk were used, against the amount of MG (pmol) was created. The MS Data were evaluated by Analyst version 1.7.2 (AB Sciex). From a linear regression the amount of MG was deduced. The limit of detection (LOD) and limit of quantification (LOQ) for the described method are 3.5 nM and 7 nM 2-MQ, respectively.

### Miscellaneous

The following experiments were performed as published before: isolation of mitochondria and *in vitro* import of radio labeled proteins (Peleh et al., 2015), fluorescence microscopy (Westermann and Neupert, 2000), and blue native gel electrophoresis (Priesnitz et al., 2020).

## Data and code availability

The mass spectrometry proteomics data (see also Tables S1 and S3) have been deposited to the ProteomeXchange Consortium via the PRIDE (Perez-Riverol et al., 2019) partner repository with the dataset identifier PXD033171 (Dynamic SILAC) and PXD033193 (Label-free data set)

Reviewer account details:

PDX033171: **Username:**

PDX033171: **Password:** KvvOw3Ww

PXD033193: **Username:**

PXD033193: **Password:** lMbku5zV

## Acknowledgments

We thank Lena Krämer, Anna Gröger, Simone Stegmüller, Sabine Knaus, Andrea Trinkaus and Ralph Mahlberg for technical assistance, Martin van der Laan, Chris Meisinger and Peter Kötter for reagents and Zuzana Storchova for support. The contribution of Katja Hansen, Naama Aviram and Maya Schuldiner was essential for the identification of Ubc8 as a factor relevant for mitochondrial protein biogenesis. This project was funded by grants from the Deutsche Forschungsgemeinschaft (HE2803/10-1 and RTG 2737 STRESSistence to JMH, and SFB1218 (project ID 269925409) to T.B.) and the Landesschwerpunkt BioComp.

## Conflict of interest

The authors declare no competing financial interests.

## Author contributions

J.M.H. conceived and supervised the study. S.R. generated and verified constructs and strains and performed the biochemical experiments. F.d.B. and T.B. analyzed data from blue native and import assays. M.R. and S.R. performed the proteomics experiments. M.R., S.R. and C.G. performed the bioinformatical analysis of the mass spectrometry data. H.B. and E.R. analyzed metabolites. J.Z., S.L. and B.M. analyzed metabolic cycles using a fermentor. J.M.H. wrote the manuscript with input from all authors.

## Supplemental Data

**Fig. S1.**
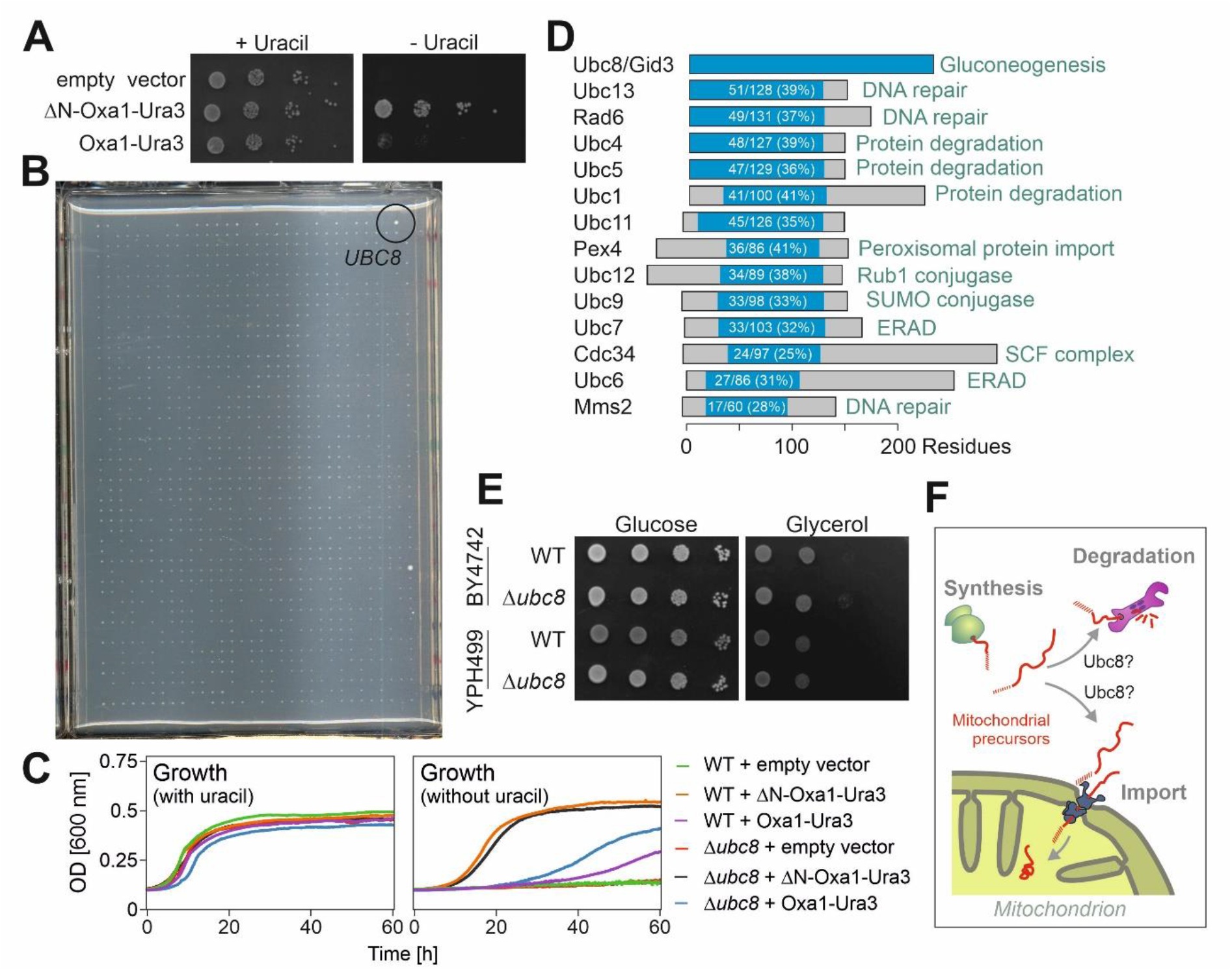
A genetic screen identified Ubc8 as a factor required for efficient depletion of mitochondrial precursors from the cytosol. **A.** Wild-type cells expressing Oxa1-Ura3 or an N-terminal truncated version of it lacking the mitochondrial targeting signal (ΔN-Oxa1-Ura3) were grown to mid-log phase. Tenfold serial dilutions were dropped onto plates containing or lacking uracil. Thus, failure in protein import results in robust growth on uracil-deficient plates. **B.** Picture of the entire uracil-deficient plate from which a sector was shown in Fig. 1B. The plate shows growth of 768 deletion mutants. Note that Δ*ubc8* and Ema35 (Laborenz et al., 2021) grew better than any other of the mutants on this plate. **C.** Growth analysis on SD medium containing or not containing uracil of the strains indicated. **D.** Graphical overview of Ubc8 homologs of yeast. **E.** The *UBC8* gene was deleted in two different genetic backgrounds and growth on glucose and the non-fermentable carbon source glycerol was tested. **F.** The accumulation of mitochondrial precursor proteins in the cytosol indicates a role of Ubc8 in their import or the proteolytic degradation.

**Fig. S2.**
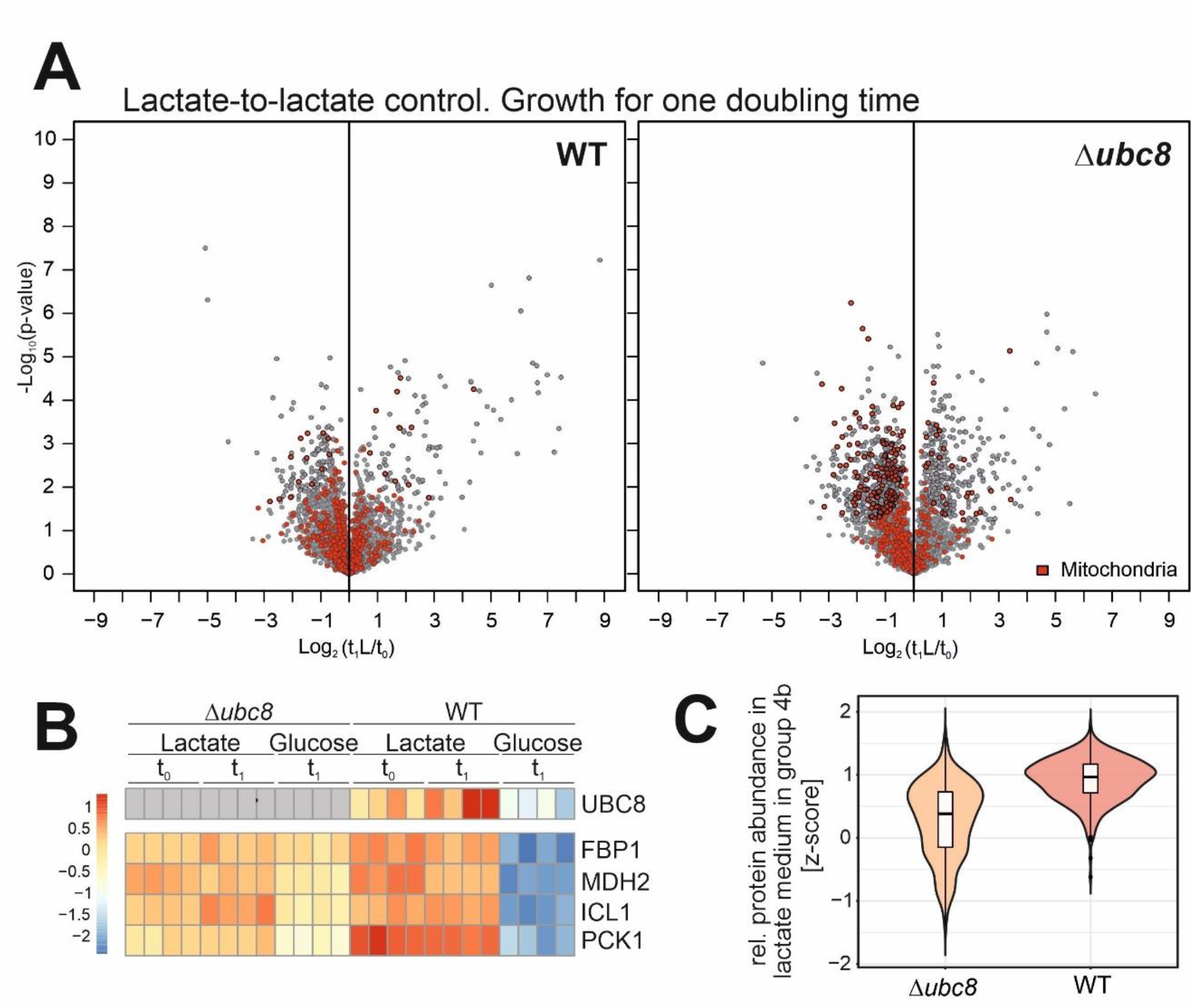
**A.** Volcano plots showing the effect of lactate-to-lactate medium replacement. In comparison to the lactate-to-glucose replacement, the changes were only very moderate. In the Δ*ubc8* cells, some mitochondrial proteins were depleted after addition of fresh medium. Names of significantly changed mitochondrial proteins are indicated. **B.** Heat map of gluconeogenesis enzymes. See legend to Fig. 2D for details. **C.** The intensities (z-scores) of all proteins of group 4b at time point t_0_ were plotted, indicating that proteins of this group were present at lower levels in the Δ*ubc8* strain compared to the wild type.

**Fig. S3.**
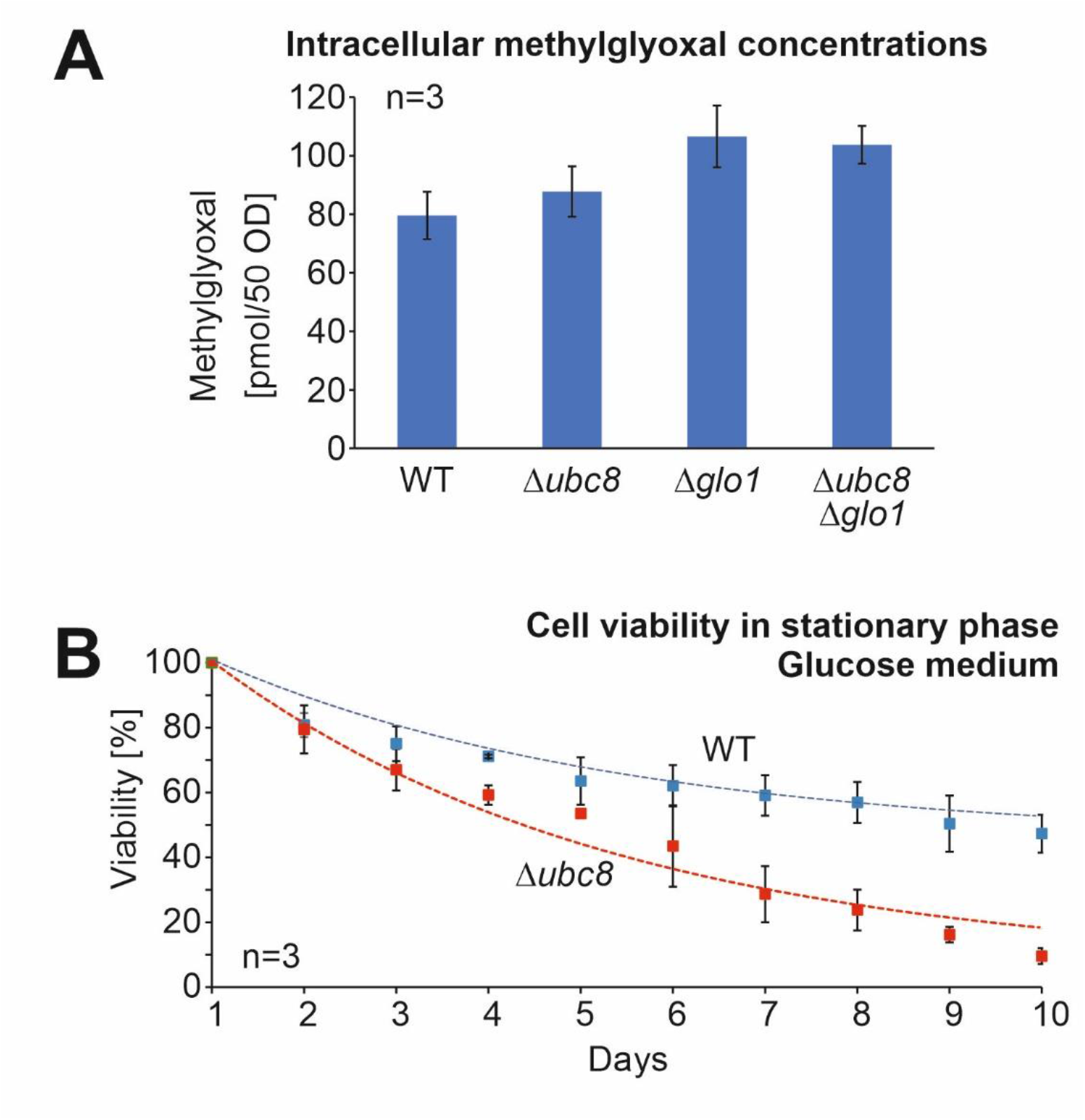
Ubc8 increases viability in stationary cultures. **A.** Yeast cells of the indicated strains were grown in 0.2% glucose to mid-log phase. Then, glucose was added to final concentration of 2%. After growth for 1 h, cells were harvested, and methylglyoxal levels were analyzed by mass spectrometry, using methylglyoxal solutions as reference. **B.** Cells were grown in 2% glucose medium to full saturation (about OD 10) and further incubated in a shaker at 30°C for 10 days. Every day, an aliquot of the culture was analyzed and viable cells counted after plating on glucose medium. Shown are mean values of three biological replicates.

**Fig. S4.**
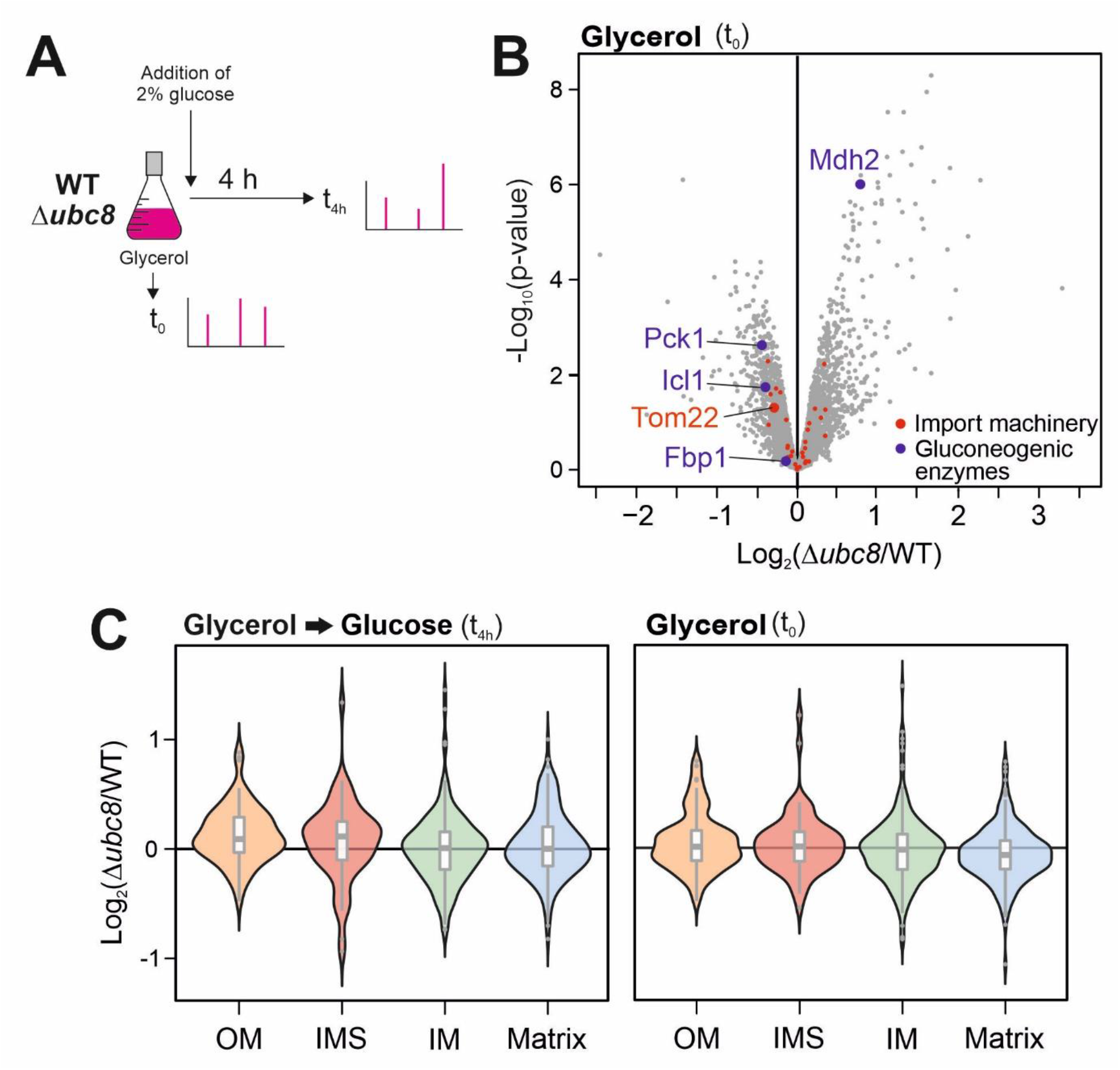
Ubc8 increases the levels of Tom22 and is particularly relevant for the accumulation of proteins of the mitochondrial matrix. **A.** Scheme of the proteome analysis. **B.** Volcano plot comparing protein levels of wild-type and Δ*ubc8* cells in glycerol-grown cultures before glucose was added. Under these conditions, gluconeogenic enzymes are not degraded, and their level is not diminished in the presence of Ubc8. The indicated proteins of the mitochondrial import machinery were Tom70, Tom71, Tom40, Tom22, Tom20, Tom6, Tom5, Mdm10, Sam50, Sam37, Sam35, Tim50, Tim23, Tim17, Tim44, Pam16, Pam18, Ssc1, Mge1, Tim21, Mgr2, Pam17, Tim22, Tim54, Sdh3, Tim8, Tim9, Tim10, Tim12, Tim13, Mas1, Mas2, Mia40 and Erv1. **C.** Violin plots to show the relative abundance of proteins of the different mitochondrial subcompartments in Δ*ubc8* cells relative to that in wild-type cells.

Supplemental Table S1. Proteomic analysis of wild-type and Δ*ubc8* cells based on dynamic SILAC labeling.

Supplemental Table S2. GO analysis for the proteins represented in groups 1, 2, 3, 4a and 4b.

Supplemental Table S3. Proteomic analysis of wild-type and Δ*ubc8* cells after shift from glycerol-induced respiration to glucose-induced fermentation for 4 h.

**Supplemental Table S4.** Antibodies used in this study.

| Type | Name | Host | Dilution | Source |
| --- | --- | --- | --- | --- |
| Primary antibody | Anti-DHFR | Rabbit | 1:1,000 | van der Laan lab |
| Primary antibody | Anti-Sod1 | Rabbit | 1:500 | Herrmann lab |
| Primary antibody | Anti-Sam50 | Rabbit | 1:50 | Becker lab |
| Primary antibody | Anti-Mdm10 | Rabbit | 1:250 | Becker lab |
| Primary antibody | Anti-Mmm1 | Rabbit | 1:500 | Becker lab |
| Primary antibody | Anti-Tom22 | Rabbit | 1:1000 | Becker lab |
| Primary antibody | Anti-Tom40 | Rabbit | 1:1000 | Becker lab |
| Primary antibody | Anti-Tom70 | Rabbit | 1:1000 | Becker lab |
| Primary antibody | Anti-Cox4 | Rabbit | 1:1000 | Becker lab |
| Primary antibody | Anti-Rip1 | Rabbit | 1:1000 | Becker lab |
| Primary antibody | Anti-Atp3 | Rabbit | 1:250 | Becker lab |
| Primary antibody | Anti-Cox1 | Rabbit | 1:1000 | Becker lab |

**Supplemental Table S5.**
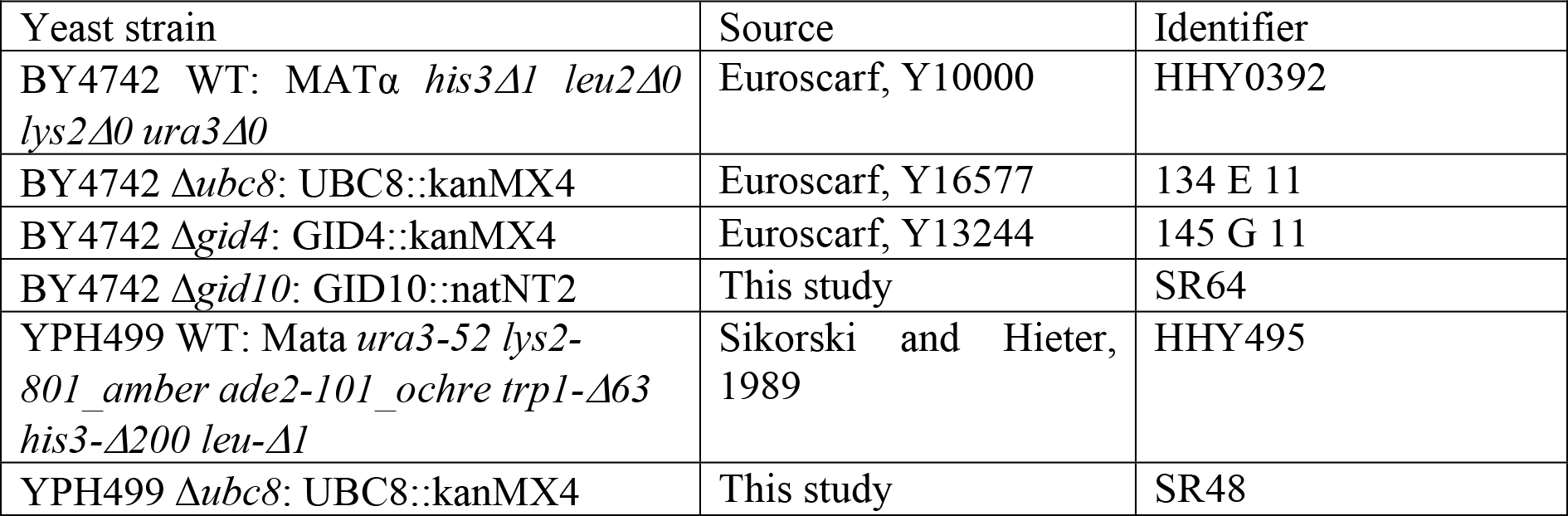
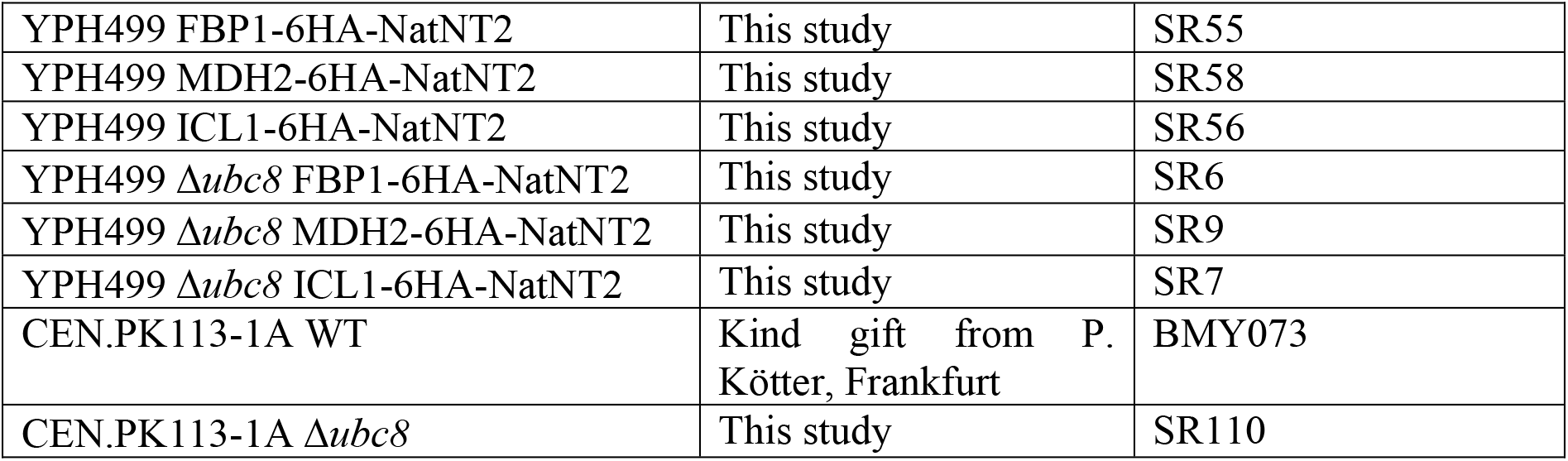
Yeast strains used in this study.

| Yeast strain | Source | Identifier |
| --- | --- | --- |
| BY4742 WT: MATα *his3Δ1 leu2Δ0 lys2Δ0 ura3Δ0* | Euroscarf, Y10000 | HHY0392 |
| BY4742 $\Delta ubc8$: UBC8::kanMX4 | Euroscarf, Y16577 | 134 E 11 |
| BY4742 $\Delta gid4$: GID4::kanMX4 | Euroscarf, Y13244 | 145 G 11 |
| BY4742 $\Delta gid10$: GID10::natNT2 | This study | SR64 |
| YPH499 WT: Mata *ura3-52 lys2-801_amber ade2-101_ochre trp1-Δ63 his3-Δ200 leu-Δ1* | Sikorski and Hieter, 1989 | HHY495 |
| YPH499 $\Delta ubc8$: UBC8::kanMX4 | This study | SR48 |
| YPH499 FBP1-6HA-NatNT2 | This study | SR55 |
| YPH499 MDH2-6HA-NatNT2 | This study | SR58 |
| YPH499 ICL1-6HA-NatNT2 | This study | SR56 |
| YPH499 $\Delta ubc8$ FBP1-6HA-NatNT2 | This study | SR6 |
| YPH499 $\Delta ubc8$ MDH2-6HA-NatNT2 | This study | SR9 |
| YPH499 $\Delta ubc8$ ICL1-6HA-NatNT2 | This study | SR7 |
| CEN.PK113-1A WT | Kind gift from P. Kötter, Frankfurt | BMY073 |
| CEN.PK113-1A $\Delta ubc8$ | This study | SR110 |

## Notes

### Competing Interest Statement

The authors have declared no competing interest.

## References

Amponsah PS, Yahya G, Zimmermann J, Mai M, Mergel S, Muhlhaus T, Storchova Z, Morgan B. 2021. Peroxiredoxins couple metabolism and cell division in an ultradian cycle. Nat Chem Biol, 17: 477–484.

Araiso Y, Imai K, Endo T. 2021. Structural snapshot of the mitochondrial protein import gate. FEBS J, 288: 5300–5310.

Araiso Y, Tsutsumi A, Qiu J, Imai K, Shiota T, Song J, Lindau C, Wenz LS, Sakaue H, Yunoki K, Kawano S, Suzuki J, Wischnewski M, Schutze C, Ariyama H, Ando T, Becker T, Lithgow T, Wiedemann N, Pfanner N, Kikkawa M, Endo T. 2019. Structure of the mitochondrial import gate reveals distinct preprotein paths. Nature, 575: 395–401.

Bausewein T, Mills DJ, Langer JD, Nitschke B, Nussberger S, Kuhlbrandt W. 2017. Cryo-EM Structure of the TOM Core Complex from Neurospora crassa. Cell, 170: 693–700 e7.

Becker T, Wenz LS, Thornton N, Stroud D, Meisinger C, Wiedemann N, Pfanner N. 2011. Biogenesis of mitochondria: dual role of Tom7 in modulating assembly of the preprotein translocase of the outer membrane. J. Mol. Biol., 405: 113–124.

Benjamini Y, Hochberg Y. 1995. Controlling the False Discovery Rate: A Practical and Powerful Approach to Multiple Testing. Journal of the Royal Statistical Society. Series B (Methodological), 57: 289–300

Bogenhagen DF, Haley JD. 2020. Pulse-chase SILAC-based analyses reveal selective oversynthesis and rapid turnover of mitochondrial protein components of respiratory complexes. J Biol Chem, 295: 2544–2554.

Boos F, Krämer L, Groh C, Jung F, Haberkant P, Stein F, Wollweber F, Gackstatter A, Zoller E, van der Laan M, Savitski MM, Benes V, Herrmann JM. 2019. Mitochondrial protein-induced stress triggers a global adaptive transcriptional programme. Nat Cell Biol, 21: 442–451.

Burnetti AJ, Aydin M, Buchler NE. 2016. Cell cycle Start is coupled to entry into the yeast metabolic cycle across diverse strains and growth rates. Mol Biol Cell, 27: 64–74.

Chacinska A, Koehler CM, Milenkovic D, Lithgow T, Pfanner N. 2009. Importing mitochondrial proteins: machineries and mechanisms. Cell, 138: 628–644.

Chen SJ, Kim L, Song HK, Varshavsky A. 2021. Aminopeptidases trim Xaa-Pro proteins, initiating their degradation by the Pro/N-degron pathway. Proc Natl Acad Sci U S A, 118.

Chen SJ, Wu X, Wadas B, Oh JH, Varshavsky A. 2017. An N-end rule pathway that recognizes proline and destroys gluconeogenic enzymes. Science, 355.

Chiang H-L, Schekman R. 1991. Regulated import and degradation of a cytosolic protein in the yeast vacuole. Nature, 350: 313–318.

Cox J, Mann M. 2008. MaxQuant enables high peptide identification rates, individualized p.p.b.-range mass accuracies and proteome-wide protein quantification. Nat Biotechnol, 26: 1367–72.

Cox J, Neuhauser N, Michalski A, Scheltema RA, Olsen JV, Mann M. 2011. Andromeda: a peptide search engine integrated into the MaxQuant environment. J Proteome Res, 10: 1794–805.

de Godoy LM, Olsen JV, Cox J, Nielsen ML, Hubner NC, Frohlich F, Walther TC, Mann M. 2008. Comprehensive mass-spectrometry-based proteome quantification of haploid versus diploid yeast. Nature, 455: 1251–4.

DeRisi JL, Iyer VR, Brown PO. 1997. Exploring the metabolic and genetic control of gene expression on a genomic scale. Science, 278: 680–6.

Di Bartolomeo F, Malina C, Campbell K, Mormino M, Fuchs J, Vorontsov E, Gustafsson CM, Nielsen J. 2020. Absolute yeast mitochondrial proteome quantification reveals trade-off between biosynthesis and energy generation during diauxic shift. Proc Natl Acad Sci U S A, 117: 7524–7535.

Dihazi H, Kessler R, Eschrich K. 2003. Glucose-induced stimulation of the Ras-cAMP pathway in yeast leads to multiple phosphorylations and activation of 6-phosphofructo-2-kinase. Biochemistry, 42: 6275–82.

Dimmer KS, Fritz S, Fuchs F, Messerschmitt M, Weinbach N, Neupert W, Westermann B. 2002. Genetic basis of mitochondrial function and morphology in Saccharomyces cerevisiae. Mol. Biol. Cell, 13: 847–853.

Doherty MK, Hammond DE, Clague MJ, Gaskell SJ, Beynon RJ. 2009. Turnover of the human proteome: determination of protein intracellular stability by dynamic SILAC. J Proteome Res, 8: 104–12.

Dong C, Zhang H, Li L, Tempel W, Loppnau P, Min J. 2018. Molecular basis of GID4-mediated recognition of degrons for the Pro/N-end rule pathway. Nat Chem Biol, 14: 466–473.

Dorrbaum AR, Kochen L, Langer JD, Schuman EM. 2018. Local and global influences on protein turnover in neurons and glia. Elife, 7.

Eilers M, Schatz G. 1986. Binding of a specific ligand inhibits import of a purified precursor protein into mitochondria. Nature, 322: 228–232.

Ellenrieder L, Opalinski L, Becker L, Kruger V, Mirus O, Straub SP, Ebell K, Flinner N, Stiller SB, Guiard B, Meisinger C, Wiedemann N, Schleiff E, Wagner R, Pfanner N, Becker T. 2016. Separating mitochondrial protein assembly and endoplasmic reticulum tethering by selective coupling of Mdm10. Nat Commun, 7: 13021.

Galdieri L, Mehrotra S, Yu S, Vancura A. 2010. Transcriptional regulation in yeast during diauxic shift and stationary phase. OMICS, 14: 629–38.

Gerbeth C, Schmidt O, Rao S, Harbauer AB, Mikropoulou D, Opalinska M, Guiard B, Pfanner N, Meisinger C. 2013. Glucose-induced regulation of protein import receptor Tom22 by cytosolic and mitochondria-bound kinases. Cell Metab, 18: 578–87.

Giaever G, Chu AM, Ni L, Connelly C, Riles L, Veronneau S, Dow S, Lucau-Danila A, Anderson K, Andre B, Arkin AP, Astromoff A, El-Bakkoury M, Bangham R, Benito R, Brachat S, Campanaro S, Curtiss M, Davis K, Deutschbauer A, Entian KD, Flaherty P, Foury F, Garfinkel DJ, Gerstein M, Gotte D, Guldener U, Hegemann JH, Hempel S, Herman Z, Jaramillo DF, Kelly DE, Kelly SL, Kotter P, LaBonte D, Lamb DC, Lan N, Liang H, Liao H, Liu L, Luo C, Lussier M, Mao R, Menard P, Ooi SL, Revuelta JL, Roberts CJ, Rose M, Ross-Macdonald P, Scherens B, Schimmack G, Shafer B, Shoemaker DD, Sookhai-Mahadeo S, Storms RK, Strathern JN, Valle G, Voet M, Volckaert G, Wang CY, Ward TR, Wilhelmy J, Winzeler EA, Yang Y, Yen G, Youngman E, Yu K, Bussey H, Boeke JD, Snyder M, Philippsen P, Davis RW, Johnston M. 2002. Functional profiling of the Saccharomyces cerevisiae genome. Nature, 418: 387–91.

Gietz D, St. Jean A, Woods RA, Schiestl RH. 1992. Improved method for high efficiency transformation of intact yeast cells. Nucl. Acids Res., 20: 1425.

Gornicka A, Bragoszewski P, Chroscicki P, Wenz LS, Schulz C, Rehling P, Chacinska A. 2014. A discrete pathway for the transfer of intermembrane space proteins across the outer membrane of mitochondria. Mol Biol Cell, 25: 3999–4009.

Hämmerle M, Bauer J, Rose M, Szallies A, Thumm M, Dusterhus S, Mecke D, Entian KD, Wolf DH. 1998. Proteins of newly isolated mutants and the amino-terminal proline are essential for ubiquitin-proteasome-catalyzed catabolite degradation of fructose-1,6-bisphosphatase of Saccharomyces cerevisiae. J Biol Chem, 273: 25000–5.

Hansen KG, Aviram N, Laborenz J, Bibi C, Meyer M, Spang A, Schuldiner M, Herrmann JM. 2018. An ER surface retrieval pathway safeguards the import of mitochondrial membrane proteins in yeast. Science, 361: 1118–1122.

Huber W, von Heydebreck A, Sultmann H, Poustka A, Vingron M. 2002. Variance stabilization applied to microarray data calibration and to the quantification of differential expression. Bioinformatics, 18 Suppl 1: S96–104.

Janke C, Magiera MM, Rathfelder N, Taxis C, Reber S, Maekawa H, Moreno-Borchart A, Doenges G, Schwob E, Schiebel E, Knop M. 2004. A versatile toolbox for PCR-based tagging of yeast genes: new fluorescent proteins, more markers and promoter substitution cassettes. Yeast, 21: 947–962.

Karayel O, Michaelis AC, Mann M, Schulman BA, Langlois CR. 2020. DIA-based systems biology approach unveils E3 ubiquitin ligase-dependent responses to a metabolic shift. Proc Natl Acad Sci U S A, 117: 32806–32815.

Keil P, Pfanner N. 1993. Insertion of MOM22 into the mitochondrial outer membrane strictly depends on surface receptors. FEBS Lett., 321: 197–200.

Kong KE, Fischer B, Meurer M, Kats I, Li Z, Ruhle F, Barry JD, Kirrmaier D, Chevyreva V, San Luis BJ, Costanzo M, Huber W, Andrews BJ, Boone C, Knop M, Khmelinskii A. 2021. Timer-based proteomic profiling of the ubiquitin-proteasome system reveals a substrate receptor of the GID ubiquitin ligase. Mol Cell, 81: 2460–2476 e11.

Kravic B, Harbauer AB, Romanello V, Simeone L, Vogtle FN, Kaiser T, Straubinger M, Huraskin D, Bottcher M, Cerqua C, Martin ED, Poveda-Huertes D, Buttgereit A, Rabalski AJ, Heuss D, Rudolf R, Friedrich O, Litchfield D, Marber M, Salviati L, Mougiakakos D, Neuhuber W, Sandri M, Meisinger C, Hashemolhosseini S. 2018. In mammalian skeletal muscle, phosphorylation of TOMM22 by protein kinase CSNK2/CK2 controls mitophagy. Autophagy, 14: 311–335.

Kulak NA, Pichler G, Paron I, Nagaraj N, Mann M. 2014. Minimal, encapsulated proteomic-sample processing applied to copy-number estimation in eukaryotic cells. Nature Methods, 11: 319–324.

Laborenz J, Bykov YS, Knoringer K, Raschle M, Filker S, Prescianotto-Baschong C, Spang A, Tatsuta T, Langer T, Storchova Z, Schuldiner M, Herrmann JM. 2021. The ER protein Ema19 facilitates the degradation of nonimported mitochondrial precursor proteins. Mol Biol Cell, 32: 664–674.

Laz TM, Pietras DF, Sherman F. 1984. Differential regulation of the duplicated isocytochrome c genes in yeast. Proc Natl Acad Sci U S A, 81: 4475–9.

Liu H, Ding J, Kohnlein K, Urban N, Ori A, Villavicencio-Lorini P, Walentek P, Klotz LO, Hollemann T, Pfirrmann T. 2020. The GID ubiquitin ligase complex is a regulator of AMPK activity and organismal lifespan. Autophagy, 16: 1618–1634.

Liu J, Barrientos A. 2013. Transcriptional regulation of yeast oxidative phosphorylation hypoxic genes by oxidative stress. Antioxid Redox Signal, 19: 1916–27.

Mathieson T, Franken H, Kosinski J, Kurzawa N, Zinn N, Sweetman G, Poeckel D, Ratnu VS, Schramm M, Becher I, Steidel M, Noh KM, Bergamini G, Beck M, Bantscheff M, Savitski MM. 2018. Systematic analysis of protein turnover in primary cells. Nat Commun, 9: 689.

Meisinger C, Rissler M, Chacinska A, Szklarz LK, Milenkovic D, Kozjak V, Schonfisch B, Lohaus C, Meyer HE, Yaffe MP, Guiard B, Wiedemann N, Pfanner N. 2004. The mitochondrial morphology protein Mdm10 functions in assembly of the preprotein translocase of the outer membrane. Dev. Cell, 7: 61–71.

Melnykov A, Chen SJ, Varshavsky A. 2019. Gid10 as an alternative N-recognin of the Pro/N-degron pathway. Proc Natl Acad Sci U S A, 116: 15914–15923.

Morgenstern M, Stiller SB, Lubbert P, Peikert CD, Dannenmaier S, Drepper F, Weill U, Hoss P, Feuerstein R, Gebert M, Bohnert M, van der Laan M, Schuldiner M, Schutze C, Oeljeklaus S, Pfanner N, Wiedemann N, Warscheid B. 2017. Definition of a High-Confidence Mitochondrial Proteome at Quantitative Scale. Cell Rep, 19: 2836–2852.

Nielsen J. 2017. Systems Biology of Metabolism. Annu Rev Biochem, 86: 245–275.

Ocampo A, Liu J, Schroeder EA, Shadel GS, Barrientos A. 2012. Mitochondrial respiratory thresholds regulate yeast chronological life span and its extension by caloric restriction. Cell Metab, 16: 55–67.

Ong SE, Blagoev B, Kratchmarova I, Kristensen DB, Steen H, Pandey A, Mann M. 2002. Stable isotope labeling by amino acids in cell culture, SILAC, as a simple and accurate approach to expression proteomics. Mol Cell Proteomics, 1: 376–86.

Peleh V, Ramesh A, Herrmann JM. 2015. Import of proteins into isolated yeast mitochondria. Methods Mol Biol, 1270: 37–50.

Perez-Riverol Y, Csordas A, Bai J, Bernal-Llinares M, Hewapathirana S, Kundu DJ, Inuganti A, Griss J, Mayer G, Eisenacher M, Perez E, Uszkoreit J, Pfeuffer J, Sachsenberg T, Yilmaz S, Tiwary S, Cox J, Audain E, Walzer M, Jarnuczak AF, Ternent T, Brazma A, Vizcaino JA. 2019. The PRIDE database and related tools and resources in 2019: improving support for quantification data. Nucleic Acids Res, 47: D442–D450.

Pratt JM, Petty J, Riba-Garcia I, Robertson DH, Gaskell SJ, Oliver SG, Beynon RJ. 2002. Dynamics of protein turnover, a missing dimension in proteomics. Mol Cell Proteomics, 1: 579–91.

Priesnitz C, Pfanner N, Becker T. 2020. Studying protein import into mitochondria. Methods Cell Biol, 155: 45–79.

Qin S, Nakajima B, Nomura M, Arfin SM. 1991. Cloning and characterization of a Saccharomyces cerevisiae gene encoding a new member of the ubiquitin-conjugating protein family. J Biol Chem, 266: 15549–54.

Qiu J, Wenz LS, Zerbes RM, Oeljeklaus S, Bohnert M, Stroud DA, Wirth C, Ellenrieder L, Thornton N, Kutik S, Wiese S, Schulze-Specking A, Zufall N, Chacinska A, Guiard B, Hunte C, Warscheid B, van der Laan M, Pfanner N, Wiedemann N, Becker T. 2013. Coupling of mitochondrial import and export translocases by receptor-mediated supercomplex formation. Cell, 154: 596–608.

Rabbani N, Thornalley PJ. 2014. Measurement of methylglyoxal by stable isotopic dilution analysis LC-MS/MS with corroborative prediction in physiological samples. Nat Protoc, 9: 1969–79.

Rao S, Schmidt O, Harbauer AB, Schonfisch B, Guiard B, Pfanner N, Meisinger C. 2012. Biogenesis of the preprotein translocase of the outer mitochondrial membrane: protein kinase A phosphorylates the precursor of Tom40 and impairs its import. Mol Biol Cell, 23: 1618–27.

Ritchie ME, Phipson B, Wu D, Hu Y, Law CW, Shi W, Smyth GK. 2015. limma powers differential expression analyses for RNA-sequencing and microarray studies. Nucleic Acids Res, 43: e47.

Ronen M, Botstein D. 2006. Transcriptional response of steady-state yeast cultures to transient perturbations in carbon source. Proc Natl Acad Sci U S A, 103: 389–94.

Sakaue H, Shiota T, Ishizaka N, Kawano S, Tamura Y, Tan KS, Imai K, Motono C, Hirokawa T, Taki K, Miyata N, Kuge O, Lithgow T, Endo T. 2019. Porin Associates with Tom22 to Regulate the Mitochondrial Protein Gate Assembly. Mol Cell, 73: 1044–1055 e8.

Saladi S, Boos F, Poglitsch M, Meyer H, Sommer F, Muhlhaus T, Schroda M, Schuldiner M, Madeo F, Herrmann JM. 2020. The NADH Dehydrogenase Nde1 Executes Cell Death after Integrating Signals from Metabolism and Proteostasis on the Mitochondrial Surface. Mol Cell, 77: 189–202 e6.

Schäfer JA, Bozkurt S, Michaelis JB, Klann K, Munch C. 2021. Global mitochondrial protein import proteomics reveal distinct regulation by translation and translocation machinery. Mol Cell.

Schmidt O, Harbauer AB, Rao S, Eyrich B, Zahedi RP, Stojanovski D, Schonfisch B, Guiard B, Sickmann A, Pfanner N, Meisinger C. 2011. Regulation of mitochondrial protein import by cytosolic kinases. Cell, 144: 227–239.

Schuldiner M, Collins SR, Thompson NJ, Denic V, Bhamidipati A, Punna T, Ihmels J, Andrews B, Boone C, Greenblatt JF, Weissman JS, Krogan NJ. 2005. Exploration of the function and organization of the yeast early secretory pathway through an epistatic miniarray profile. Cell, 123: 507–19.

Schüle T, Rose M, Entian KD, Thumm M, Wolf DH. 2000. Ubc8p functions in catabolite degradation of fructose-1, 6-bisphosphatase in yeast. EMBO J, 19: 2161–7.

Sherpa D, Chrustowicz J, Qiao S, Langlois CR, Hehl LA, Gottemukkala KV, Hansen FM, Karayel O, von Gronau S, Prabu JR, Mann M, Alpi AF, Schulman BA. 2021. GID E3 ligase supramolecular chelate assembly configures multipronged ubiquitin targeting of an oligomeric metabolic enzyme. Mol Cell, 81: 2445–2459 e13.

Shiota T, Imai K, Qiu J, Hewitt VL, Tan K, Shen HH, Sakiyama N, Fukasawa Y, Hayat S, Kamiya M, Elofsson A, Tomii K, Horton P, Wiedemann N, Pfanner N, Lithgow T, Endo T. 2015. Molecular architecture of the active mitochondrial protein gate. Science, 349: 1544–8.

Shiota T, Mabuchi H, Tanaka-Yamano S, Yamano K, Endo T. 2011. In vivo protein-interaction mapping of a mitochondrial translocator protein Tom22 at work. Proc. Natl. Acad. Sci. USA, 108: 15179–15183.

Sikorski RS, Hieter P. 1989. A system of shuttle vectors and host strains designed for efficient manipulation of DNA in Saccharomyces cerevisiae. Genetics, 122: 19–27.

Slavov N, Macinskas J, Caudy A, Botstein D. 2011. Metabolic cycling without cell division cycling in respiring yeast. Proc Natl Acad Sci U S A, 108: 19090–5.

Smets B, Ghillebert R, De Snijder P, Binda M, Swinnen E, De Virgilio C, Winderickx J. 2010. Life in the midst of scarcity: adaptations to nutrient availability in Saccharomyces cerevisiae. Curr Genet, 56: 1–32.

Tsuboi T, Viana MP, Xu F, Yu J, Chanchani R, Arceo XG, Tutucci E, Choi J, Chen YS, Singer RH, Rafelski SM, Zid BM. 2020. Mitochondrial volume fraction and translation duration impact mitochondrial mRNA localization and protein synthesis. Elife, 9.

Tu BP, Kudlicki A, Rowicka M, McKnight SL. 2005. Logic of the yeast metabolic cycle: temporal compartmentalization of cellular processes. Science, 310: 1152–8.

Tucker K, Park E. 2019. Cryo-EM structure of the mitochondrial protein-import channel TOM complex at near-atomic resolution. Nat Struct Mol Biol, 26: 1158–1166.

Tyanova S, Temu T, Cox J. 2016. The MaxQuant computational platform for mass spectrometry-based shotgun proteomics. Nat Protoc, 11: 2301–2319.

van Wilpe S, Ryan MT, Hill K, Maarse AC, Meisinger C, Brix J, Dekker JT, Moczko M, Wagner R, Meijer M, Guiard B, Hönlinger A, Pfanner N. 1999. Tom22 is a multifunctional organizer of the mitochondrial preprotein translocase. Nature, 401: 485–489.

Wallace DC. 2005. A mitochondrial paradigm of metabolic and degenerative diseases, aging, and cancer: a dawn for evolutionary medicine. Annu. Rev. Genet., 39: 359–407.

Wang W, Chen X, Zhang L, Yi J, Ma Q, Yin J, Zhuo W, Gu J, Yang M. 2020. Atomic structure of human TOM core complex. Cell Discov, 6: 67.

Westermann B, Neupert W. 2000. Mitochondria-targeted green fluorescent proteins: convenient tools for the study of organelle biogenesis in Saccaromyces cerevisiae. Yeast, 16: 1421–1427.

Williams CC, Jan CH, Weissman JS. 2014. Targeting and plasticity of mitochondrial proteins revealed by proximity-specific ribosome profiling. Science, 346: 748–51.

Wu X, Li L, Jiang H. 2018. Mitochondrial inner-membrane protease Yme1 degrades outer-membrane proteins Tom22 and Om45. J Cell Biol, 217: 139–149.

Zeng Y, Pan Q, Wang X, Li D, Lin Y, Man F, Xiao F, Guo L. 2019. Impaired Mitochondrial Fusion and Oxidative Phosphorylation Triggered by High Glucose Is Mediated by Tom22 in Endothelial Cells. Oxid Med Cell Longev, 2019: 4508762.

